# Single-cell morphodynamical trajectories enable prediction of gene expression accompanying cell state change

**DOI:** 10.1101/2024.01.18.576248

**Authors:** Jeremy Copperman, Ian C. Mclean, Sean M. Gross, Jalim Singh, Vaibhav Murthy, Young Hwan Chang, Alexander E. Davies, Daniel M. Zuckerman, Laura M. Heiser

## Abstract

Extracellular signals induce changes to molecular programs that modulate multiple cellular phenotypes, including proliferation, motility, and differentiation status. The connection between dynamically adapting phenotypic states and the molecular programs that define them is not well understood. Here we develop data-driven models of single-cell phenotypic responses to extracellular stimuli by linking gene transcription levels to “morphodynamics” – changes in cell morphology and motility observable in time-lapse image data. We adopt a dynamics-first view of cell state by grouping single-cell trajectories into states with shared morphodynamic responses. The single-cell trajectories enable development of a first-of-its-kind computational approach to map live-cell dynamics to snapshot gene transcript levels, which we term MMIST, Molecular and Morphodynamics-Integrated Single-cell Trajectories. The key conceptual advance of MMIST is that cell behavior can be quantified based on dynamically defined states and that extracellular signals change the overall distribution of cell states by altering rates of switching between states. We find a cell state landscape that is bound by epithelial and mesenchymal endpoints, with distinct sequences of epithelial to mesenchymal transition (EMT) and mesenchymal to epithelial transition (MET) intermediates. The analysis yields predictions for gene expression changes consistent with curated EMT gene sets and predicts expression of thousands of RNA transcripts through extracellular signal-induced EMT and MET with near-continuous time resolution. The MMIST framework leverages true single-cell dynamical behavior to generate molecular-level omics inferences and is broadly applicable to other biological domains, time-lapse imaging approaches and molecular snapshot data.

**Summary:** In normal homeostatic tissues, extracellular signals induce changes in the behavior and state of epithelial cells, and aberrant responses to such signals are associated with diseases. To decode and potentially steer these responses, it is essential to link live-cell behavior to molecular programs; however, high-throughput molecular techniques are destructive or require fixation. Here we present a novel computational approach to connect single-cell measures of cell phenotype and behavior to bulk molecular readouts, enabling prediction of dynamic changes in gene expression programs. This reveals molecular programs associated with distinct cell states and identifies drivers that may be manipulated to control cell state change.

## Introduction

Cells continuously sense and respond to cues from their local microenvironment, activating molecular programs that drive changes in cell state. This exquisite control of cell state is critical for normal tissue function, and dysregulation of cell state control is increasingly recognized as a disease mechanism^1–4^. Cell state is determined by molecular and cellular composition, including genome and chromatin structure^5,6^, proteomic^7^ and transcriptomic levels^8^, mitochondrial function^9^, and metabolic activity^10^. Cell state is intrinsically mutable^11^ and is influenced by various physical^12–14^, and chemical^15,16^ cues. Single-cell omic analyses have provided an unprecedented catalog of cell states across both normal and diseased tissues^17,18^ while spatially-resolved sequencing^19^ and highly multiplexed imaging^20–22^ have revealed insights into their spatial organization; however, these approaches lack single-cell time-ordered information, limiting the ability to draw mechanistic insights. Live-cell imaging, on the other hand, readily captures cellular dynamics over timescales of seconds to days but is limited to a small number of molecular read-outs^11,23–25^. Further, analysis of live-cell data typically relies on single timepoint “snapshots” of cell morphology or fluorescently-labeled reporters^26–29^. To overcome these limitations, we recently developed a morphodynamical trajectory embedding method that leverages hidden information from time-ordered live-cell trajectories, enabling improved prediction of future behavior and improved extraction of causal and mechanistic relationships as compared to single-snapshot analyses^30^.

It is increasingly appreciated that mechanistic understanding of both normal and diseased biological systems will require consideration of cell state dynamics. Protein biomarkers have long been used to identify cell states and associated gene expression patterns^31,32^. Several recent methods impose a dynamical model upon static single-cell measurements to describe gene expression dynamics^33–35^, including pseudo-time estimation^36,37^ and RNA velocity^38,39^. In contrast, here we develop our computational framework based upon direct observation of single-cell dynamics obtained from live-cell imaging^34–37^. Broad efforts have been undertaken to map cell morphology and motility to gene expression states^26,40–44^, including generative machine learning approaches^45,46^. We take a dynamics-oriented approach, where we resolve a cell state landscape over hundreds of “microstates”, where transitions among microstates are described in a discrete-time Markov model framework^47,48^. Our data-driven modeling approach, Molecular and Morphodynamics-Integrated Single-cell Trajectories (MMIST), extends other efforts based on live-cell imaging and trajectory analysis^49,50^ by characterizing cell state changes quantitatively observed in live-cell imaging data, yielding distinct states that can be linked to molecular programs observed in companion profiling data. The central and novel element of MMIST is a mapping between morphodynamical “states” defined using live-cell features measured over time and companion bulk RNAseq data. The mapping exploits a linear population-matching approach^51–54^ based upon the premise that live-cell imaging and bulk molecular profiling share commonly identifiable cell states with shared average transcript levels specific to each state. In this framework, treatment conditions induce changes in state-switching rates leading to differences in state populations, which are directly measured from live-cell imaging. Given this shared set of states across ligand-treatment conditions, state-specific transcription levels are readily computed in a linear algebra framework.

We developed our approach by focusing on the well-characterized human mammary epithelial MCF10A cell line^55,56^, which recapitulates key features of epithelial biology, including migration^57,58^ and organoid formation^59,60^. It is also easily manipulated in a variety of assays including live-cell imaging^61^, knock-down^56^, and chemical perturbation^62^ and therefore is commonly used for cell biology studies. Prior studies have used MCF10A cells to probe epithelial responses to growth factors and cytokines^63^ and to uncover molecular programs associated with EMT^64–70^.

We examined cellular and molecular responses to soluble ligands known to alter cell state of epithelial cells--including Transforming Growth Factor Beta (TGFB) as an illustrative example— and demonstrate the quantitative linkage of EMT-associated live-cell phenotypic responses with EMT molecular programs that we validated in an independent dataset^71^. In total, our novel data-driven modeling approach captures cell state change along sequences of cell state intermediates via live-cell and gene expression phenotypes and enables linkage of imaging and molecular data to uncover molecular correlates of distinct morphodynamic cell states.

## Results

### Approach to integrate morphodynamical and gene expression measurements to inform cell state change

Our method is designed to infer molecular programs associated with distinct cell states by linking morphodynamic measurements acquired in live-cell imaging data to companion snapshot molecular data. We analyze a recently published LINCS MCF10A ligand perturbation dataset^63^ that consists of paired live-cell imaging and bulk transcriptomic measurements of MCF10A cells after treatment with the ligands Epidermal Growth Factor (EGF), Transforming Growth Factor Beta (TGFB), and Oncostatin M (OSM).

Our computational framework, illustrated in **Figure 1**, leverages companion live-cell image stacks and bulk RNAseq as input, and utilizes statistical physics approaches to yield maps of cell states and their transition sequences^72,73^. Here we outline the major steps. (a) First, we analyze the live-cell image data to identify cell nuclei by training a virtual nuclear reporter^74^ on paired phase contrast and nuclear reporter images, then virtually stain nuclei in the entire dataset (**Supplementary** Figure 1). We initially “featurize” individual cells to quantify cell shape and texture and perform local environment featurization using geometric boundaries based on the nuclear centers. We assessed the qualitative performance of the MMIST pipeline using cytoplasmic masks obtained using cellpose software on a subset of the data (**Supplementary** Figure 2). We track individual cells across images with Bayesian belief propagation^75^ and compute motility as cell displacement between frames. Local alignment of cell motility is featurized via a neighborhood averaging approach (**Supplementary** Figure 3). Assessment of segmentation and tracking performance is provided in **Supplementary Data Table 1**. (b) Dynamical cell features are constructed based on trajectory snippets (all possible single-cell sub-trajectories of a particular length in a sliding window manner, e.g., frames 1-2-3, 2-3-4, 3-4-5, …) utilizing our morphodynamical trajectory embedding methodology^30^. Thus, the features used for analysis are morphodynamical trajectory snippets, quantified as time-ordered lists of morphological features. (c) Morphodynamical trajectory snippets are used to build a data-driven dynamical model of cell states. (d) Cell states observed in the image data are mapped to gene transcript levels using linear decomposition. This yields temporal sequences of morphodynamical cell state changes and their associated gene expression levels.

**Figure 1:**
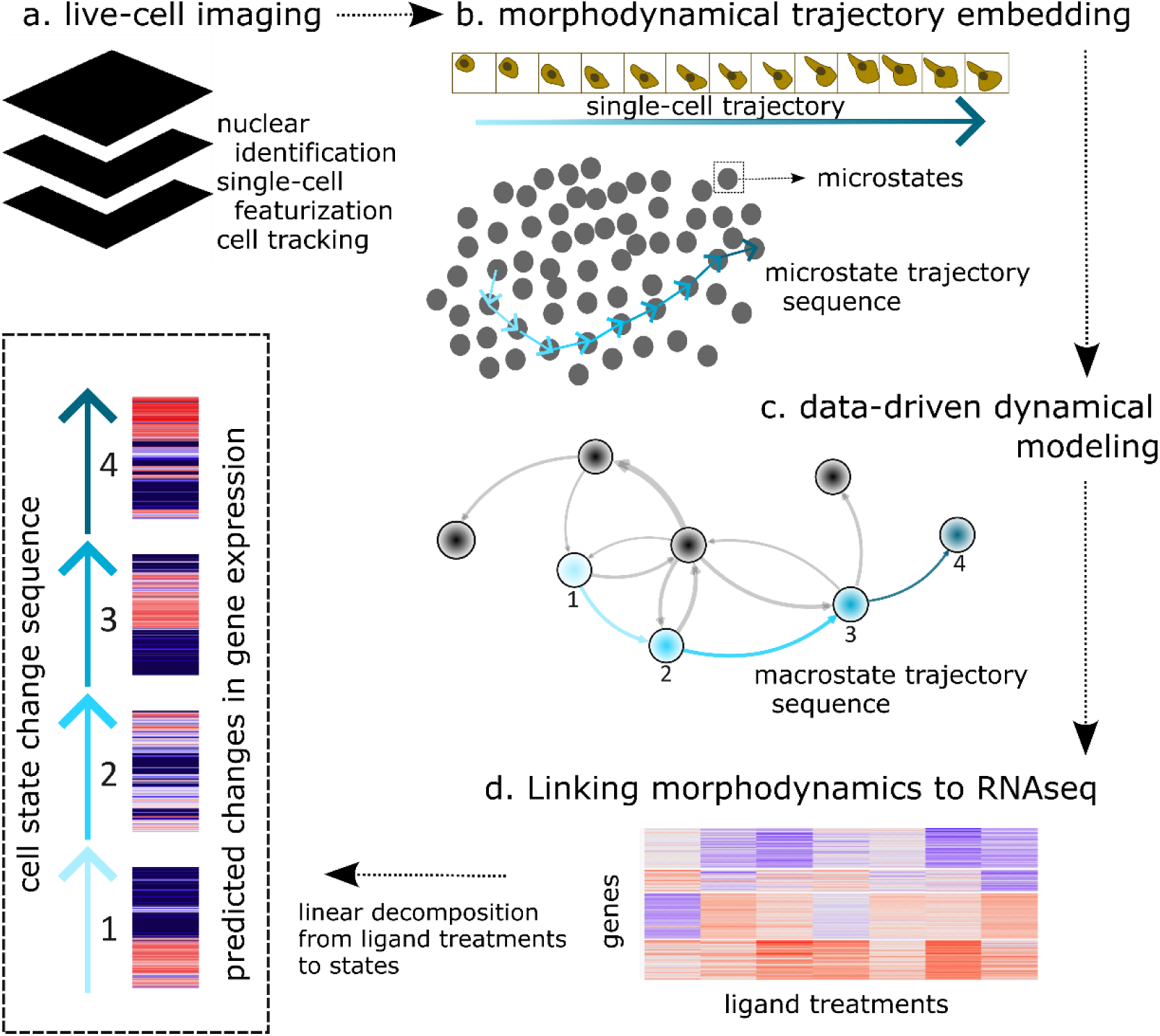
MMIST approach to link live-cell imaging to molecular read-outs. a) Live-cell imaging of MCF10A cells after treatment with a panel of microenvironmental ligands. Nuclei are identified using a convolutional neural network, and single-cells are featurized and tracked through time. b) Single-cell features are concatenated along single-cell trajectories to construct the morphodynamical trajectory embedding. c) Dynamical models learn cell states and cell state change sequences in the morphodynamical landscape. d) Cell state populations are used as a linear decomposition of bulk gene expression measurements to predict the gene expression programs underlying cell state change.

### Single-cell trajectories define morphodynamical cell states

Morphodynamical states form the basis of our analysis. We define these states as groups of cells that exhibit similar trajectories in shape, texture, and motility over time. To identify morphodynamic states, we employ feature vectors that are a time-concatenation of image-based features^30^; this is termed the ‘morphodynamical trajectory space.’ In this space, we place hundreds of “microstate” centers via clustering. We then count transitions among microstates to build a data-driven transition matrix Markov model of cell state progression^47,48^. Next, microstates are grouped into coarser “macrostates” using a spectral clustering procedure^76,77^.

We refer to these macrostates as morphodynamic states or simply states. The eigenfunctions of the dynamical model represent dynamical motifs, which we visualize using UMAP^78^ dimensionality reduction to facilitate interpretation of cell states (**Figure 2a**). Clustering the microstates revealed 14 discrete morphodynamic states, each with characteristic features, notably with the formation of multicellular clusters as a key feature driving this partitioning (**Figure 2b**. We examined the relationship between morphodynamic states and ligand treatment and found that ligands induce unique cell state changes and transition flows between macrostates, indicating induction of different cell state populations over time in response to perturbation (**Figure 2c**). For example, OSM drives transition flow towards state 10, consisting of highly motile multicellular clusters, whereas TGFB drives transition flow towards state 5, consisting of more isolated cells with extended lamellopodia-like cytoskeletal features. The complete set of ligand-dependent cell state populations are shown in **Supplementary** Figure 4c, and population distributions and transition flows are shown in **Supplementary** Figure 5.

**Figure 2:**
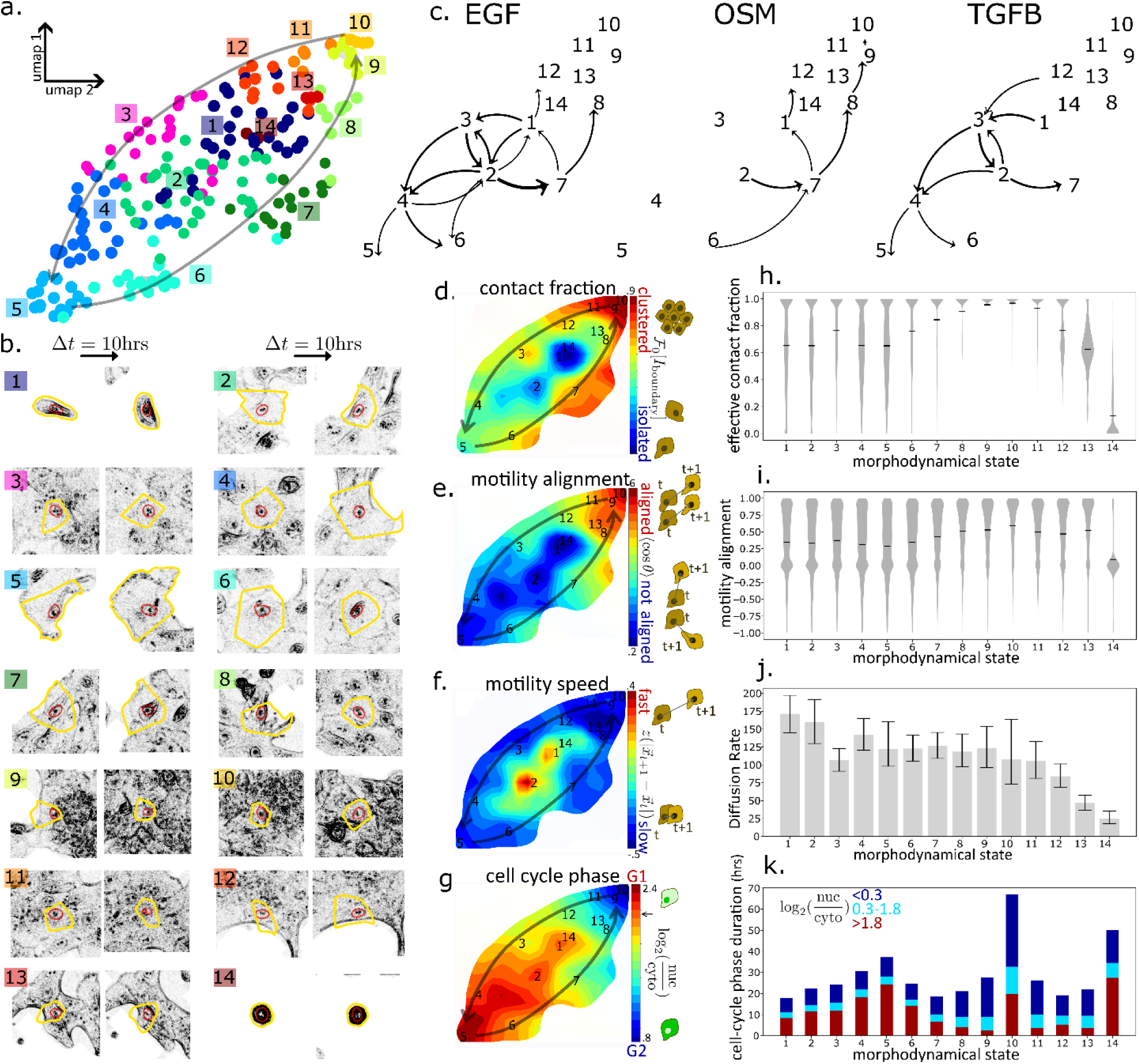
Data-driven models define morphodynamical cell states and state transition dynamics. a.) The dynamical embedding landscape is visualized via UMAP from 200 microstates (dots) constructed from morphodynamical trajectories (trajectory snippet length = 10H), large black arrows are guides to the eye for significant macroscopic flow paths. b.) Images from first and last frames of representative trajectory snippets (10H trajectory length) from each state with nuclear segmentations (red contours) and associated Voronoi segmentation (yellow contours). c.) Cell state flow (at t=24H) by ligand treatment. d.-g.) Cell morphology, motility, and cell cycle features by morphodynamical cell state. Panels (h) and (i) show violin-plot distributions of single-cell values, (j) shows average diffusion rate (pixels^2^/timestep) with uncertainty based on single-cell variation, and (k) shows modeled cell-cycle phase durations averaged over single-cell behavior.

The derived states resolve differences in morphodynamical properties, including cell-cell contact fraction, local alignment of cell-cell motility, motility speed, and cell-cycle phase (**Figure 2d-g**). We quantified these morphological properties for each cell state to highlight state-to-state differences (**Figure 2h-k**). States 5 and 10 at the extremes of the morphodynamical cell state space. State 5 is characterized by mesenchymal-associated features such as lower local alignment of cell motility, more extended cytoskeletal features, greater cell spreading, and an extended G1 cell cycle duration (**Figure 2k**); this state population increases under TGFB containing treatments. In contrast, state 10 harbors epithelial-associated features, including increased multicellular clustering and collective motility, which are increased after treatments that include OSM. Intermediate states between states 5 and 10 have fewer cell-cell contacts (**Figure 2d,h**), increased motility (**Figure 2f,j**), and short cell cycle duration (**Figure 2g,k**). Under EGF treatment, cells transition between these intermediate states (**Figure 2c**). Overall, the morphodynamically-derived cell state space matches the well-described biological framework of epithelial and mesenchymal cell states^79^, including extended G1 duration in the mesenchymal state^80–83^. Below, we discuss these observations in the context of EMT.

### Cross-modal mapping of morphodynamic trajectories reveals temporal variation in molecular responses

Motivated by the observation that morphodynamic cell states recapitulate aspects of EMT, we next sought to identify molecular programs associated with cell state transitions. This process relies on having both morphodynamical observations and molecular measurements for an identical set of experimental treatments. The primary assumption is that an observed morphodynamical state corresponds to similar RNA levels regardless of ligand treatment. Consider the example of linking RNA levels to two distinct states (motile and non-motile), where the cell state frequencies are modulated by ligand treatment. If ligand A induces an increase in the motile cell state population as compared to ligand B and *also* induces higher RNA levels for gene X, then we infer that motility is linked to expression of gene X. This qualitative idea can be made exact in a simple linear algebra framework by decomposing each measured average transcript level as a linear sum over morphodynamical state populations and gene expression profiles (**Supplementary** Figure 4c).

We validated the linear population matching approach by assessing its capability to predict withheld gene expression levels in ligand combination conditions. Specifically, we withheld the OSM+EGF, EGF+TGFB, and the triple combination OSM+EGF+TGFB RNA-seq data from the training set so that we could predict gene-expression profiles for these three conditions. This approach requires the same number of states as training conditions (ligand treatments), so we performed a new clustering into 10 morphodynamical cell states, using the remaining 11 treatments as training data and averaging over the 11 possible decompositions over the 10 states. Because the morphodynamical state space is unchanged, these 10 states span the same morphodynamical features as the 14 states described in Figure 2, just more coarsely. The morphodynamical cell state populations from the live-cell imaging in the withheld test set treatments, combined with morphodynamical cell state decomposed gene expression levels from the training set, enable prediction of the RNA-seq in the test set. Overall, the measured and predicted global gene expression levels show similar patterns (**Figure 3a**). We assessed the predictive capability of the model at multiple trajectory lengths by computing the Pearson correlation between measured and predicted differential gene expression for 13,516 genes (**Supplementary Data Table 2**). Prediction quality is maximal at a trajectory length of 10 hours (r > 0.7, p-value<0.001, upper-tailed test). The predictions out-perform >99% of null model-predictions based on randomized state populations (**Figure 3b**). As a linear decomposition-based method, overall correlation between MMIST predictions and measured gene expression is limited by the capability of the training treatment gene expression to span the withheld test treatment gene expression. To test how robust MMIST was to training and test set splits, we tested MMIST in a leave-one-out k-fold cross validation setting; MMIST performance significantly exceeded null prediction in all training/test splits and could predict the gene expression better than any closest training set treatments in 5/6 test folds (**Supplementary** Figure 6). As a further test of MMIST-predicted gene expression dynamics, we analyzed companion cyclic immunofluorescence time-course data available on these samples^63^. We compared expression dynamics of cognate RNA-protein pairs, which revealed MMIST-predicted changes in RNA levels over time correlated with measured protein expression over time for most gene targets, at a similar scale to the direct correlation between protein and RNA levels and far exceeding a null expectation with predictions randomly scrambled over time (**Supplementary** Figure 7). These findings provide support for the validity of our approach to link morphodynamical states observed in image data to companion molecular measurements.

**Figure 3:**
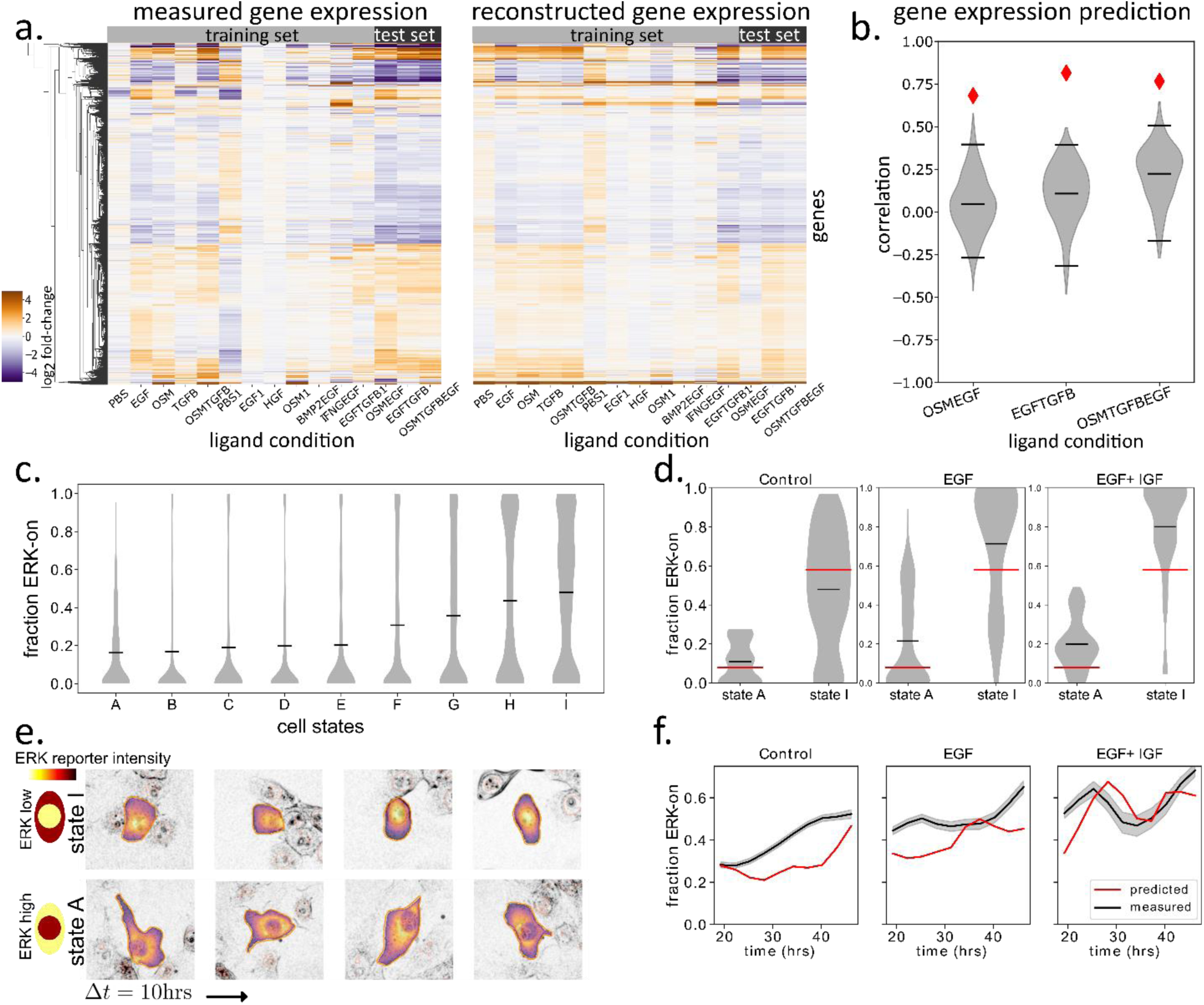
Morphodynamical cell states predict global gene expression patterns and live-cell ERK reporter activity. a.) Validation of model gene expression predictions: measured and model-reconstructed gene expression at 24hrs for every experimental condition, including training set (light gray) and test set conditions. b.) Correlation between measured and model-predicted gene expression (red diamonds), and null estimates using random state populations (gray violin plots). Horizontal lines are the mean, 5^th^, and 95^th^ percentile of the null distribution. c.) Fraction of ERK “on” cells (ERKTR c/n ratio >0.8) (violin plots, mean as black line) by morphodynamical cell state. d.) Fraction of ERK on cells within highest and lowest ERK activity morphodynamical cell states by ligand condition (violin plots, mean as black lines) and MMIST derived morphodynamical state level (red lines). e.) Representative cell images in morphodynamical cell states, brightfield in black and ERKTR color-mapped within cell segmentation boundary. f.) Fraction of ERK “on” cells over time by treatment, measured mean and bootstrapped 95% CI (black line and shaded area), and predicted using MMIST (red line).

In a final validation, we assessed the ability of MMIST to predict temporal changes in molecular responses using experimental approaches that enable quantitative real-time measures of molecular activity, facilitating evaluation of our approach. Specifically, we genetically engineered malignant basal-like breast cancer cell line T4-2 cells to express an ERK translocation reporter^11,84^. Cells were treated with a panel of 12 ligand treatments including EGF, IGF, and the combination EGF+IGF then subjected to brightfield and fluorescence imaging every 15 minutes for 48 hours. We applied our morphodynamical cell state analysis pipeline with a trajectory length of 16 hours, extracting 9 morphodynamical cell states. As expected, the predicted morphodynamical cell states showed variable ERK activity (**Figure 3C** and **Supplementary** Figure 8). We binarized ERK activity into “on” and “off” states and measured the fraction of ERK “on” cells in the window between 24-48 hours. For a direct single-cell validated comparison in the morphodynamical cell states, we calculated the fraction or ERK “on” states along each single-cell trajectory of 10 hours. We identified morphodynamical cell states associated with low and high ERK activity under both control and ligand-stimulated conditions (**Figure 3d,e**), supporting our hypothesis that morphodynamic cell states can also be defined by similar pathway activity across ligand treatments, and morphodynamic cell state population shifts can aggregate to yield an overall population-level response to perturbation. From the live-imaging extracted morphodynamical state populations and the snapshot bulk measured ERK active cell fraction at 24 hours, MMIST predicted the dynamical progression of ERK active cell fractions over the imaging duration (**Figure 3f**). In this independent cell line and molecular readout, MMIST successfully predicted live-reporter derived pathway activity, confirming MMIST capability to link molecular and morphodynamic cell phenotypes.

### MMIST identifies ligand-induced EMT and MET morphodynamical cell state change sequences

As an illustrative use case of MMIST, we next focused on morphological features associated with canonical epithelial and mesenchymal cell states. Analysis of the morphodynamical cell states revealed features canonically associated with epithelial and mesenchymal cell states, including changes in cell-cell motility alignment and cell clustering, as described above (**Figure 2d-j**). Motivated by prior studies of epithelial-mesenchymal transition (EMT) and mesenchymal-epithelial transition (MET)^33,50,66,71^, we used our framework to examine the relationship between these end states and the influence of microenvironmental signals in mediating transitions between them. For EMT, we assigned state 1 as the most highly populated state at the initial trajectory time window (10 hours), while state 5 was assigned as mesenchymal due to its mesenchymal-like morphological features and enrichment in the TGFB condition. For MET, we assign state 8 as the final state because it is enriched over time in the EGF treatment as cells reach confluence (**Supplementary** Figure 9), and the initial state is assigned as state 5, the endpoint of our EMT sequence.

We observe robust but distinct state transition sequences for the EMT and MET transitions, consistent with a highly driven nonequilibrium system^85,86^. For EMT the dominant sequence of states is (1,2,3,4,5) while for MET, the primary sequence is (5,6,7,8), shown in **Figure 4a,b**. Transitions back to state 1 are common in most treatments (**Figure 2b** and **Supplementary** Figure 5). The dominant sequences of state changes are robust across ligand treatments, though the probability of specific state-to-state transitions varies. For instance, OSM treatment drives most cells towards dense and collectively migrating epithelial-like clusters (state 10), but for the rare cells that do reach state 5 from state 1, the dominant sequence of states remains the same (**Supplementary** Figure 10).

**Figure 4:**
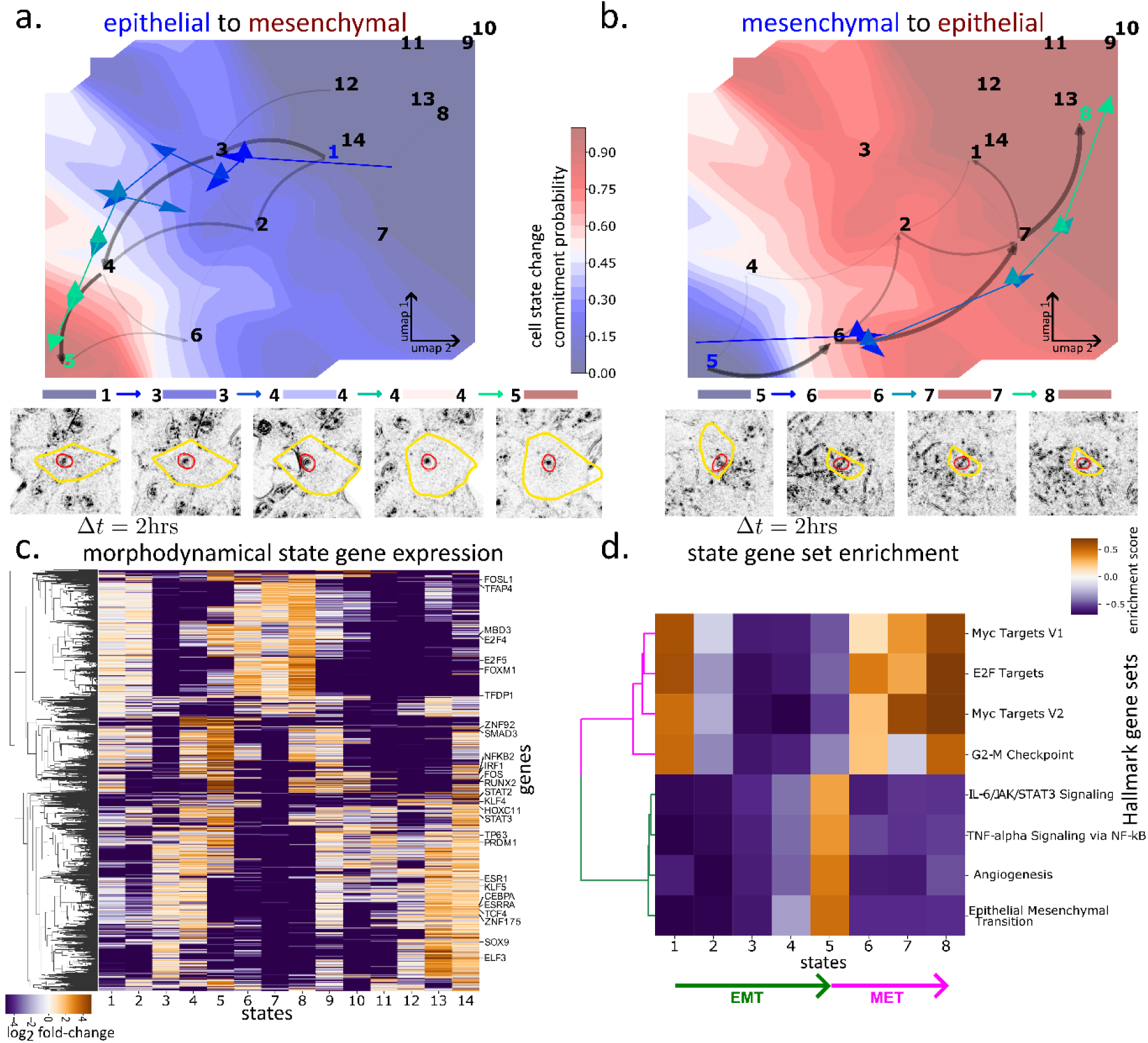
Concerted gene expression changes along EMT/MET morphodynamical cell state change sequences. a.-b.) Cell state change pathways (black arrows; thickness proportional to probability flux carried by each state-to-state transition) based on cell states from Figure 2a, and cell state change commitment probability (blue to red) in EGF (reference positive control) condition. Also shown are representative single-cell trajectory (dark blue to turquoise arrows, 30min timestep) and cell images (2 hrs between images). c.) Differential gene expression in each morphodynamical cell state (top 8000 most variable genes), and transcription factors identified in ligand response molecular modules from Gross et al. ^63^ labeled on y-axis. d.) Hallmark gene set enrichment over EMT/MET associated cell states.

MMIST revealed unique expression patterns associated with each morphodynamical cell state (**Figure 4c**). We performed gene set enrichment over the Hallmark gene sets^87^ on the derived morphodynamical state gene expression profiles. The morphodynamical state-decomposed gene expression along the EMT state change sequence shows a transition from a proliferative program enriched for Hallmark Myc Targets V1 and V2, E2F targets, and G2-M transition, to a mesenchymal program enriched for IL4/JAK/STAT3, TNFA via NFKB, Angiogenesis, and Epithelial to Mesenchymal Transition (**Figure 4d**). This switch from a proliferative program to a mesenchymal gene expression program augments our observation that cell-cycle phase durations co-vary with mesenchymal-like features observed in the live-cell data (**Figure 2j**).

### Near-continuous gene expression time evolution prediction during TGFB-driven EMT

MMIST yields near-continuous time evolution of morphodynamical cell state populations via a Markov model calibrated by counting transitions between microstates extracted from single-cell trajectories. For example, EGF+TGFB leads to an increase in mesenchymal-like states 4, 5, and 6, whereas these states are decreased after EGF-only treatment (**Figure 5b**). We assessed model predictions by comparing state population proportions measured in live-cell imaging as a function of time after EGF+TGFB treatment. The model recapitulates the morphodynamical state population trends observed in the live-cell imaging data (**Figure 5b**). While quantitative differences in the timescale of population changes indicate that stationarity and the Markov assumption are only partially satisfied, overall trends are well captured, notably decreases in the populations of states 7 and 8, and increases in states 4, 5, and 6. Final state populations (48 h) predicted from the 30 m-lag Markov model are accurate to within 2% absolute probability for all states.

**Figure 5:**
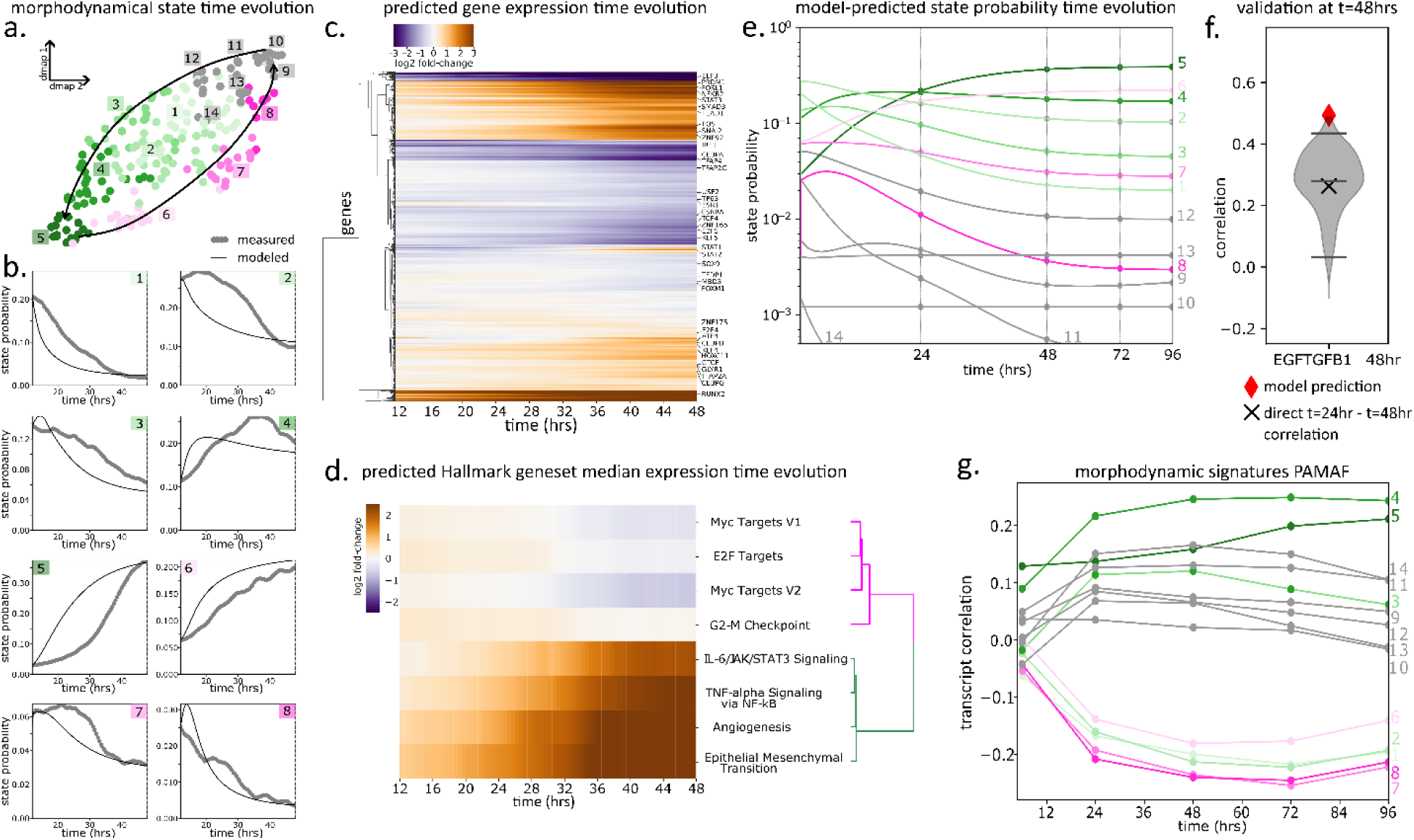
Morphodynamical model predicts EGF+TGFB-induced EMT gene expression time evolution. When predicted RNA levels of morphodynamic states are integrated with Markov model dynamics, an array of dynamical omics predictions can be made, shown here for the EGF+TGFB condition. a) Morphodynamical states, which are numbered 1-12 and color-coded (mesenchymal: green, epithelial: purple). Color labels for the states are consistent throughout figure. b) State probability time evolution, measured (grey dots) and model-derived (black lines), with y-axis limits set for each plot so small changes in state populations are visible. c) Prediction of gene expression over time at 30-minute intervals using morphodynamical state prediction and live-cell imaging measured state probabilities, with transcription factors identified from Gross et al. ^63^, and d) summarized to Hallmark gene sets. e) Model-predicted state probability time evolution over 96 hours, trained from live-cell imaging over 48 hours. f) Correlation between measured and model-predicted gene expression at t=48H (red diamond) based on training data from t=24H, relative to null models with random state probabilities (gray distribution). Also shown: correlation between t=24H and t=48H gene expression (black X). g) Correlation between predicted morphodynamical state gene signatures and PAMAF measurements out to 4 days.

Our computational framework also enables a prediction of gene transcript levels at the same near-continuous time intervals (dt=30min) as those measured in the live-cell image data. Conceptually, we make these predictions by first associating each morphodynamical state with a full gene expression profile (including all genes measured in RNAseq) and then predicting the bulk gene expression over time by computing the weighted sum of the gene expression profiles for states observed in each treatment condition (**Figure 5c**). Under EGF+TGFB, our model predicts a continuous shift in multiple gene programs, including decreases in proliferation-associated programs and increases in mesenchymal-associated programs (**Figure 5d**). These changes are quantified based on average expression over the set of genes contained in the Hallmark gene sets^87^ with statistically significant enrichment in the morphodynamical state decomposition analysis (**Figure 4d**)

MMIST can also be used to predict future, unmeasured shifts in cell state populations. For example, the model predicts large shifts in state populations between 0-48H, which we observed experimentally; however, it also predicts continued subtle shifts in state populations beyond the 48H duration of the experiment (**Figure 5e**). We next assessed the ability of our model to predict unseen changes in gene expression programs. Here, we trained our model with RNA-seq data collected at 24H post-treatment, then used it to predict gene expression profiles at 48H based on the predicted morphodynamical state populations shown in **Figure 5e**. We assessed our predictions by computing the correlation between experimentally measured and predicted expression profiles, after normalizing to t=0H. MMIST-predicted gene expression profiles at t=48H exhibit a Pearson correlation of ∼0.5 with withheld 48H RNA-seq data, outperforming the ∼0.25 correlation between 24H and 48H experimental profiles (**Figure 5f** and **Supplementary** Figure 4). This indicates that MMIST predictions can capture molecular programs associated with morphodynamic state change.

The transcriptional programs associated with TGFB-driven EMT have been previously investigated in MCF10A cells, and datasets generated through these efforts provide a useful tool for independent validation of our model^66,67,70^. To evaluate the EMT-associated signature extracted via our morphodynamical analysis, we compare our results to a recently published, independent, time-resolved gene expression dataset of MCF10A cells treated with EGF+TGFB then harvested for molecular profiling at multiple timepoints including 24, 48, 72, and 96 hours post-treatment, (“PAMAF” data)^71^; this dataset lacked companion live-cell image data. We first assessed the biological significance of the model-assigned morphodynamic states based on gene expression levels, finding positive correlation between PAMAF measurements and mesenchymal-like morphodynamical cell states 4 and 5 after EGF+TGFB treatment (**Figure 5g**). Consistent with this, epithelial-like states 6, 7, and 8 are among the least correlated. Together, these findings provide support for the robustness of MMIST to identify meaningful biological signals that can be validated in independent data sets.

## Discussion and Conclusion

The novelty of MMIST lies in the ability to derive morphology/motility-based cell states which may co-exist in various single-cell ligand treatments, and to map these cell states to a unique gene expression profile. No broadly available experimental approach enables time-resolved molecular profiles with a similar level of detail. Multimodal integration with single-cell molecular measurements, such as single-cell RNAseq, provide increased opportunities for future development. Our approach provides a direct map from live-cell derived sequences of morphodynamical cell state changes to comprehensive gene expression profiles. Using unlabeled imaging, MMIST-derived cell states predicted ligand response gene expression dynamics, provided mechanistic insight into the sequence of intermediates along EMT/MET, and was validated at the single-cell level and population level in an independent live-reporter cell line.

We explored a paradigm in which individual cells transition between morphodynamic states with treatment-specific dynamics and state frequencies. The resultant trajectory space captures states induced by different perturbations, where these perturbations alter cell state transition rates. This report demonstrates the value of this paradigm, as it enables mapping of complex, spatiotemporal phenotypes to gene transcript levels. One limitation of the present model is that it is restricted to the range of behaviors observed for a particular cell type (MCF10A) under the treatments examined and does not represent a comprehensive assessment of all possible cell states. Thus, the derived (coarse-grained) dynamical models are incomplete. As live-cell information increases, for instance via the incorporation of multiplexed live-cell reporters and deep-learning based image featurization^42,43,88–90^, integration with fixed single-cell and spatial omics profiling at endpoint may require a separation of shared information across cell populations from unique information to each single cell^91^.

From a physical theoretical point of view, the transition mechanism of a dynamical process is defined via the set of trajectories connecting two states of interest^72,92–94^, for instance epithelial and mesenchymal cell states. The single-cell trajectory set that connects these end states contains the set of intermediate transition states^49^. Here, we have captured sequences of EMT and MET intermediates, consistent with the emerging view of epithelial and mesenchymal states as a continuum^17,33^. An open question is whether characterization of transition intermediates will yield insight into cell state control, which could inform the control of EMT-driven processes during development or disease progression, such as tumor invasion^95,96^. Future studies could extend our findings by employing inhibitor or gene knockout approaches to functionally assess EMT transition intermediates predicted to be critical for cell state control.

Cell state biomarkers can predict sensitivity to targeted drugs^2,97^, and are expressed in a spatially organized manner in both healthy and diseased tissues^98,99^. Morphodynamical cell state definitions can expand upon known biomarker-based cell states, providing a prediction of the dynamical responses to biological manipulation. We expect that linking morphodynamics to gene expression changes, in spatial context, will lead to a deeper understanding and ability to control cell state changes in complex tissue and tissue-like environments.

Characterization of the transition mechanism via live-cell image-based trajectories, such as we have presented, is not a mechanistic explanation at the molecular level. Time-ordered single-cell trajectories of the quantity of molecular species, such as gene transcripts, imply but do not prove causality. Molecularly detailed single-cell trajectory data to constrain mechanistic models can provide predictions of causal molecular relationships that could be experimentally validated. Similarly, our data-driven approach does not yield a prediction for unmeasured perturbations, for instance response to different ligands or drugs. We speculate that mechanistic models^100–103^, trained using the type of detailed trajectory data at the molecular level we have presented here, may enable prediction of cell behavior in unseen contexts.

Live-cell phenotypic responses to ligand perturbation are well-described by our single-cell morphodynamical trajectory-based data-driven modeling approach and enable a mapping between live-cell phenotype and time-dependent gene expression changes. Our models yielded a validated prediction of near-continuous gene expression levels during ligand-driven EMT/MET in MCF10A cells. MMIST is broadly applicable across biological contexts, perturbations, and molecular read-outs. We envision that it will be useful characterize and inform cell state dynamics in developmental, homeostatic, and disease settings.

## Methods

### MCF10A Cell Culture

**C**ompanion RNAseq and live-cell imaging data used in this paper are described in full in Gross, et al^63^. In brief, MCF10A cells were cultured in growth media composed of DMEM/F12 (Invitrogen #11330-032), 5% horse serum (Sigma #H1138), 20 ng/ml EGF (R&D Systems #236-EG), 10 µg/ml insulin (Sigma #I9278), 100 ng/ml cholera toxin (Sigma #C8052), 0.5 µg/ml hydrocortisone (Sigma #H-4001), and 1% Pen/Strep (Invitrogen #15070-063). For all ligand response experiments, cells were seeded in growth media in collagen-coated well plates and allowed to attach for 6-hours. Cells were then washed with PBS, and growth media was replaced with growth-factor free media lacking EGF and insulin. After an 18-hour incubation, cells were treated with ligands in fresh growth-factor free media. Seven ligand conditions were tested at concentrations determined to elicit maximal cell responses^63^(10 ng/ml EGF (R&D Systems #236-EG), 40 ng/ml HGF (R&D Systems #294-HG), 10 ng/ml OSM (R&D Systems #8475-OM), 20 ng/ml BMP2 (R&D Systems #355-BM) + 10 ng/ml EGF, 20 ng/ml IFNу (R&D Systems #258-IF) + 10 ng/ml EGF, 10 ng/ml TGFβ (R&D Systems #240-B) + 10 ng/ml EGF). Cells were then subjected to live-cell imaging (imaged every 30 minutes for 48 hours), followed by bulk RNAseq. Here these samples are specified by appending a “1” to the treatment condition (e.g. EGF1) in order to distinguish these treatments from the cell cycle reporter studies.

### Cell cycle reporter studies

To assess cell-cycle responses to ligand treatments, MCF10A cells were genetically modified to stably express the HDHB cell-cycle reporter^104^ and a red nuclear reporter. These MCF10A cells were a gift from Gordon B. Mills and from the same aliquot as the LINCS MCF10A dataset described above^63^. The methodology used to generate the reporter cell line has been described previously^105^. Reporter cells were treated with ligand (EGF 10 ng/ml (R&D Systems #236-EG), OSM 10 ng/ml (R&D Systems #8475-OM), TGFB 10 ng/ml (R&D Systems #240-B), EGF 10 ng/ml + OSM 10 ng/ml, TGFB 10 ng/ml + EGF 10 ng/ml, OSM 10 ng/ml + TGFB 10 ng/ml, TGFB 10 ng/ml + EGF 10 ng/ml + OSM 10 ng/ml)and then imaged every 15 minutes for 48 hours with an Incucyte S3 microscope (1020x1280, 1.49 𝜇𝑚/pixel). Three channels were collected – phase contrast, red (nuclear) and green (cell-cycle) – for four fields of view per well. The initial frame coincided with the addition of the ligands and fresh imaging media. ^63^

### RNA-seq analysis

Detailed description of sample preparation, processing, and alignment can be found in Gross et al^63^. For each ligand treatment, we performed a differential expression analysis from time 0H controls on the RNA-seq gene-level summaries with the R package DESeq2 (1.24.0), with shrunken log2 fold change estimates calculated using the apeglm method. We applied a minimum expression filter such that log2(TPM)>0.5 in at least 3 measurements over treatments and replicates (with TPM transcripts per million), yielding 13,516 genes with measured differential expression from control used in our analysis.

### Image preprocessing

Foreground (cells) and background pixel classification was performed using manually trained random forest classifiers using the ilastik software^106^. Images were z-normalized (mean subtracted and normalized by standard deviation). In cell images, absolute values of these z-normalized pixel values are shown (white to black). Image stacks were registered translationally using the pystackreg implementation of subpixel registration^107^.

### Nuclear segmentation

A convolutional neural network was trained to predict the nuclear reporter intensity from the matched phase contrast images for imaging data of WT MCF10A cells with no nuclear reporter. In the EGF, OSM, and TGFB conditions, 4 image stacks (12 total) were used to train the FNET 3D reporter prediction convolutional neural network CNN from the Allen Cell Science Institute^74^, with time as the third dimension. This trained CNN was then used to predict nuclear reporter channel from the bright-field image over all image stacks in datasets. See **Supplementary** Figure 1 for representative nuclear reporter prediction and comparison to ground truth. Nuclear segmentations were generated by performing a local thresholding of the image within 51 pixel-sized windows at 1 standard deviation of intensity. Segmentations were filtered for a minimum size of 25 pixels and a maximum size of 900 pixels, see **Supplementary Table 1** for segmentation performance. To capture features including the local environment around a single nucleus, the image was partitioned into Voronoi cells around each nuclear center, with background classified pixels removed.

### Cell featurization

Single-cell featurization was performed on the Voronoi-partitioning of the image by nuclear center. Cell features are described in detail in Copperman et al.^30^ and repeated here for convenience. Morphology features were obtained as follows: segmented cells were extracted, and mask-centered into zero-padded equal sized arrays larger than the linear dimension of the biggest cell (in each treatment). Principal components of each cell were aligned, and then single-cell features were calculated. Zernike moments (49 features) and Haralick texture features (13 features) were calculated in the Mahotas^108^ image analysis package. The sum average Haralick texture feature was discarded due to normalization concerns. Rotation-invariant shape features (15 features) were calculated as the absolute value of the Fourier transform of the distance to the boundary as a function of the radial angle around cell center^109^, with the set of shape features normalized to 1. The cell environment was featurized in a related fashion. First, an indicator function was assigned to the cell boundary with value 0 if the boundary was in contact with the background mask, and value 1 if in contact with the cell foreground mask. The absolute value of the Fourier transform of this indicator as a function of radial angle around cell-center then featurized the local cell environment (15 features), with the sum of cell environment features normalized to 1. Note the first component of the cell environment features is practically the fraction of the cell boundary in cell-cell contact. The high-dimensional cell feature space was dimensionally reduced using principal component analysis (PCA), retaining the largest 11 eigen-components of the feature covariance matrix (spanning all treatments and image stacks) which captured >99% of the variability. PCA features primarily captured cell shape and local cell-cell contact information (**Supplementary** Figure 11a).

### Motility features

Cell motility was characterized in a single-cell manner, referenced both to the image frame and relative to neighboring cells. Single-cell displacement 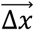 between tracked frames was z-normalized, and cells which could not be tracked backward for a frame had unrecorded displacements and were not used in our analysis. The local motility alignment of a single-cell to the local neighborhood of contacting cells (sharing a Voronoi boundary) was measured by extracting the cosine of the angle between the single-cell and direct neighbors via 𝑝^_1_ · 𝑝^_2_ with 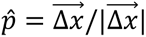. Local contact inhibition of locomotion was measured via the higher-order vector formed by (𝑝^_1_ − 𝑝^_2_) · 𝑟^_12_ with 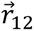 the separation vector between cells^110^. Neighborhood averages were taken via the Voronoi partition, averaged over neighbors and weighted by the relative length of the boundary to each neighbor, see **Supplementary** Figure 3.

### Batch normalization

The live-cell imaging data analyzed were collected from two separate imaging studies. Single-cell featurization can depend in subtle ways upon the imaging treatment and sample batch. To normalize these effects we utilized a batch normalization procedure at the single-cell feature level. For each morphology feature, we utilized a histogram matching procedure between negative control (PBS) treatments. We then fit a linear model to the histogram-matched distributions, and applied this linear model between sample batches, see **Supplementary** Figure 10b.

### Cell tracking

To follow single-cells through time to extract the set of single-cell trajectories for morphodynamical trajectory embedding, we utilized a Bayesian likelihood-based approach implemented in the btrack software package^75^ using default parameters. This Bayesian approach was applied for each frame over a 12 frame window, and then successful tracks over each pair of successive frames were extracted. See **Supplementary Table 1** for manual validation of tracking performance.

### Morphodynamical trajectory embedding

To maximize the single-cell information, we extended single-timepoint morphology and motility features over single-cell trajectories using a delay-embedding approach, described in Copperman et al.^30^ In brief, single-cell features including motility features, but excluding cell-cycle features, were concatenated along the trajectory length to form morphodynamical feature trajectories. We tested multiple trajectory lengths and selected a trajectory length of 10 hours where the best prediction of withheld treatment combination RNA-seq was obtained, see **Supplementary Table 2**. We utilized a dynamical embedding approach described below to cluster trajectories and visualize this space, and did not perform any further dimensionality reduction upon the trajectory concatenated morphological feature PCAs and motility feature trajectories prior to dynamical model building.

### Data-driven dynamical Markov state model

To capture dynamical properties within the morphodynamical space, we constructed a transition matrix Markov model within the trajectory embedding space. The embedded space was binned into “microbins” using k-means clustering with 𝑘 = 200 clusters. As we collected millions of single-cell transitions, sampling exceeds common Markov model heuristics regarding transition sampling^47^, and results using 50, 100, 200, and 400 clusters are qualitatively similar. In this discrete space, a transition matrix 𝑇 between bins was estimated from the set of transition counts 𝐶_𝑖𝑗_ from microbin *i* to *j* as 𝑇_𝑖𝑗_ = 𝐶_𝑖𝑗_/𝐶_𝑖_ with 𝐶_𝑖_ = ∑_𝑗_ 𝐶_𝑖𝑗_. The matrix 𝑇 represents transition probabilities for a lag time of 30 minutes. This accounting was agnostic to cell birth and death processes, yet we observe our model reproduces the main trends of morphodynamical state population evolution, see Figure 5b. Coarse-graining of the Markov model is described below: see “Cell state change pathways.”

### Dynamical features

To evaluate live-cell behavior via characterization of shared dynamics, we have applied a dynamical featurization approach via the data-driven transition matrix model. Using a transition matrix model constructed from all possible single-cell trajectory steps in the in the microbinned trajectory feature space, we construct the Hermitian extension 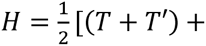 𝑖(𝑇 − 𝑇′)] with 𝑇′ the transpose of the transition matrix 𝑇, this approach numerically stabilizes the eigendecomposition and provides all real eigenvalues for unambiguous ordering of eigencomponents^111^. We retain 15 dominant eigencomponents (see **Supplementary** Figure 12), and concatenate real and imaginary parts of eigenvectors to construct a 30-dimensional characterization of each microbin center. To visualize the dynamical trajectory space, we apply UMAP dimensionality reduction of the microstate eigenvector components to 2 components. Average flows in the UMAP space are calculated via calculating microstate dependent average displacements via the transition matrix < 𝑥_𝑖_ >= ∑_𝑗_(𝑥_𝑖_ − 𝑥_𝑗_)𝑇_𝑖𝑗_ and averaging over 10 nearest microstate neighbors for smoothness. We note that UMAP flows were used only for visualization, not featurization.

### Morphodynamical cell states

As a tool for reducing complexity and extracting biological meaning in the morphodynamical embedding space, we defined a set of macrostates by clustering together microstates using dynamical similarity. Specifically, we utilize the eigencomponents of 𝐻 (Hermitian extension of the transition matrix 𝑇, see Dynamical Embedding) and perform k-means in the kinetic motifs. We utilize a lower cutoff of 0.015 for the total fraction of cell trajectories assigned to each state; if a microstate has too few trajectories assigned, then it is combined with its nearest neighbor by Euclidean distance in the space of dynamical motifs. The k-means clusters are increased until the requested number of macrostates with minimum fraction assignment is obtained. We then evaluated the capability of the derived macrostates to describe the state-change dynamics by evaluating the sum of timescales captured in the macrostate transition matrix model, related to the VAMP score^112^. We observe a rapid increase in score increasing to 10 states and continued increase beyond 15 states, see **Supplementary** Figure 12. Note that the macrostates, like the features themselves, were not designed or optimized for the task of predicting RNA levels.

### Cell state change pathways

To extract the sequences of morphodynamical cell states under EMT/MET, we adopted a transition path approach to calculate commitors and state change sequences utilizing our data-driven Markov model^72^. Fine-grained transition matrices were coarse-grained by morphodynamical cell state (macrostate) membership, i.e., by summing over transition counts from associated microstates based on the same lag time (30 min), and flux analysis was carried out using the PyEMMA analysis package^73^; all pathways carrying flux between sets of initial and final states were evaluated to find dominant state change sequences. Committor probabilities (for reaching the final state before returning to the initial state) were highly dependent upon culture treatment, but cell state change sequences were quite robust to culture treatment, see **Supplementary** Figure 10.

### Cell-cycle reporter analysis and dynamical modeling

To capture cell-cycle dynamics from the HDHB reporter images, we adopted a similar data-driven modeling approach as we took in defining the morphodynamical cell states. Reporter levels in the nuclear and cytoplasmic compartments were extracted, and the ratio of these reporter levels was used as a self-normalizing readout of cell-cycle state, where exclusion of HDHB from the nucleus is known to correlate with G2 cell-cycle state, with maximal nuclear correlation occurring abruptly at mitosis and decreasing gradually from G1 to S, and with minimal nuclear signal at G2^105^. To divide reporter ratio values into cell-cycle stages, we utilized our Markov state modeling and dynamical embedding procedure, first building a microbin model with 50 bins evenly spaced throughout the range of reporter ratio values, then dividing these into 4 macrostates via k-means clustering in dynamical motifs, see **Supplementary** Figure 13. We then calculated mean first passage times using PyEMMA between cell-cycle stages as a readout of cell-cycle stage lifetimes in each of the morphodynamical cell states.

### Bulk RNAseq reconstruction

To capture the biological drivers of morphodynamical cell state changes, we mapped our morphodynamical cell states (defined above) to RNA-seq-based gene expression profiles. We adopted a linear decomposition approach based on the single assumption that the identical set of morphodynamic states occurs in all experimental conditions so that each state exhibits the same average gene profile in all conditions. In this framework, if cells in treatment A are subdivided into a set of states 𝑠 with known state populations 𝑝*_s_*^𝐴^ such that ∑*_s_* 𝑝*_s_*^𝐴^ = 1, and the state and treatment dependent average gene levels are known, a bulk measurement of the 𝑖th gene can be reconstructed exactly as 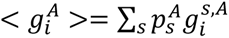, where 𝑔*_i_*^𝑠,𝐴^ is the average expression level for gene *i* in state *s* under experimental condition A. We approximate this exact expression by making the assumption that cells in state 𝑠 under each treatment have identical *average* gene expression, i.e., that 𝑔*_i_*^𝑠,𝐴^ = 𝑔*_i_*^𝑠^ regardless of A, for every 𝑠. The utility of this approximation can be evaluated via our results, and is equivalent to letting the states form a non-negative matrix factorization of the bulk expression. Under this assumption, we have a linear system of equations connecting state populations and state gene expression levels 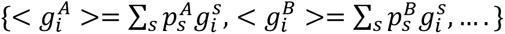, one equation for each treatment A,B,C,… based on treatment-specific cell state populations 𝑝*_s_*^𝐴^, 𝑝*_s_*^𝐵^, … directly measured via live-cell imaging and morphodynamical analysis, and with paired bulk RNA-seq measurements < 𝑔*_i_*^𝐴^ >. If there are as many measurements as states, this linear equation can be inverted for the gene expression profiles in each state, 𝑔*_i_*^𝑠^. If there are less states than treatments and the solution is over-determined, we obtain the solution over all possible combinations of treatments and average over the results. In practice, state definitions and gene expression measurements contain errors, and true solution of the linear system would yield negative gene levels, so we do a least squares minimization with the constraint of positive gene levels. We use fold-changes rather than absolute gene levels to preserve the batch and replicate normalization, this normalization does not affect the system of equations as it enters on both sides of the equality. To validate our state decomposition of measured bulk RNA-seq pipeline, we split our data into training sets and validation sets. State gene expression levels are trained from the training set gene levels only, and gene expression for withheld test set conditions are then predicted via the measured morphodynamical cell state populations. Null model predictions are constructed from uniformly distributed random state populations combined with previously estimated state-specific gene levels (from true populations) as a measure of how unique the measured state populations are at predicting the test set gene expression.

### Gene set enrichment

To interpret morphodynamical cell state gene expression profiles, we performed gene set enrichment analysis via the pyGSEA package^113^. We utilized the preranked algorithm, sorting genes via the predicted gene expression levels in each morphodynamical cell state. We ran gene set enrichment using the Hallmark gene sets^87^, which broadly capture well-studied biological processes and cell signaling activity.

### Live-cell ERK activity imaging and analysis

To validate MMIST predictions from our MCF10A cell line data, we leveraged an independent mammary epithelial cell line that has been used to quantify heterogeneity in ERK activity and downstream gene expression^11^. HMT-3522 cell lines were cultured in DMEM/F-12 (Gibco, Cat# 11320033) supplemented with 1.4×10⁻⁶M hydrocortisone (Sigma-Aldrich, Cat# H4001-1G), 10⁻¹⁰M β-estrogen (Sigma-Aldrich, Cat# E2758-1G), 2.6ng/mL sodium selenite (Sigma-Aldrich, Cat# S5261-100G), 10 µg/mL transferrin (Sigma-Aldrich, Cat# T8158-1G) and 250ng/mL insulin (Sigma-Aldrich, Cat# I1882-100MG) as previously described^114^. For live imaging experiments, reporter cell lines were generated via lentiviral transduction of ERK and AKT translocation reporter fused to mVenus and mStrawberry respectively, in addition to nuclear reporter H2B fused to iRFP670. Cells were sequentially selected with puromycin, blasticidin and neomycin to ensure the stable co-expression of all three reporters.

Live-cell imaging was performed using 96-well plates with #1.5 glass bottoms (Cellvis, P96-1.5 H-N), pre-coated with 25 µl of 50 µg/mL rat tail collagen Type I (ThermoFisher, A1048301) in 20 mM acetic acid for 1 hour. The plates were washed with PBS to remove excess and unbound collagen. T4-2 reporter cells were seeded at a density of 21,750 cells per well and allowed to adhere overnight. The following morning, cells were rinsed twice with phenol-free DMEM/F-12 medium (Gibco, 21041-025) without additives and subsequently placed in imaging media containing phenol-free DMEM/F-12 supplemented with 1.4×10⁻⁶ M hydrocortisone, 10⁻¹⁰ M β-estrogen, and 10 µg/mL transferrin and allowed to reach steady state for 4 hours prior to imaging.

Imaging was performed using a Nikon Ti2 automated microscope equipped with an Oko-Lab environmental chamber set to 37 °C and 5% CO₂, a SOLA II LED illumination system, and a Photometrics Prime 95B camera. The system was operated via NIS-Elements AR software. Images from each well were captured every 15 minutes for 48 hours using a Nikon Plan Apo 20X air objective with a numerical aperture of 0.80. Baseline ERK phosphorylation was assessed during an initial imaging period of four hours, after which treatments were applied sequentially: the first treatment (T1) for two hours followed by the second treatment (T2). The treatment order was as follows: Control (T1), EGF (T1), IGF (T1), EGF+IGF (T1), MEKi (T1), MEKi+EGF (T2), MEKi+IGF (T2), EGF+MEKi (T2), IGF+MEKi (T2), HGF (T1), AKTi (T1), AKTi+EGF (T2), and EGF+AKTi (T2). Ligands and inhibitors used in the experiment include EGF (100ng/mL, Peprotech Cat# AF-100-15-500UG), IGF (100ng/mL, Peprotech Cat# 100-11-500UG), HGF (100ng/mL, Peprotech Cat# 100-39H-25UG), MEKi/PD0325901 (500nM, Selleckchem Cat# S1036), AKTi/MK2206 (1µM, Selleckchem Cat# S1078).

Cytoplasmic and nuclear regions were segmented using cellpose software^115^, tracked between frames, with brightfield-derived single-cell morphology and motility features calculated as described above with our celltraj software. ERK translocation reporter (ERKTR) activity was quantified as the ratio of the mean ERKTR intensity in cytoplasmic to nuclear regions, in each cell. ERK activity was binarized into ERK off and ERK on states at a cutoff of ERK c/n=0.8, roughly 1 standard deviation above the mean ERK c/n ratio under inhibition treatment (MEKi). The fraction ERK on was calculated as the fraction of cells with ERK c/n ratio above the cutoff in the entire treatment population, and as the fraction of time a single cell had an ERK c/n ratio above the cutoff along the single-cell trajectory. We performed our trajectory embedding analysis at a trajectory length of 16 hours, roughly the autocorrelation decay time of the single-cell features. We extracted 9 morphodynamical cell states, and the MMIST pipeline was applied taking the population fraction of ERK on cells at t=24 hrs as the bulk molecular measurement which we decomposed into morphodynamical cell state mean activity levels, see **Supplementary** Figure 8. ERK activity dynamics were predicted in the MMIST framework using the extracted morphodynamical cell state populations from the live imaging. Further details can be found in the analysis script maintained in the celltraj software repository.

## Acknowledgements

We thank Joe Gray for contributions to project conceptualization, Mark Dane for technical guidance and assistance accessing LINCS data, David Aristoff and Gideon Simpson for mathematically oriented discussions, and John Russo and Luke Ternes for input regarding computational implementation. J.C. was supported by the Damon Runyon Cancer Research Foundation Quantitative Biology Fellowship DRQ-09-20. Y.H.C is supported in part by the National Cancer Institute (U54CA209988, U2CCA233280). D.M.Z. acknowledges support from the National Science Foundation (MCB 2119837). L.M.H. acknowledges support from NIH research grants U54CA209988 and U54HG008100, and the Anna Fuller Foundation. The authors acknowledge Lauren Kronebusch for help with manuscript editing.

## Data and Code Availability

All codes and scripts to perform the analysis in this work can be found at the project github repository.

https://github.com/jcopperm/celltraj

LINCS MCF10A Molecular Deep Dive data is available in some formats from the synapse database^63^ and additional data is available upon request.

---

**Supplementary Figure 1:**
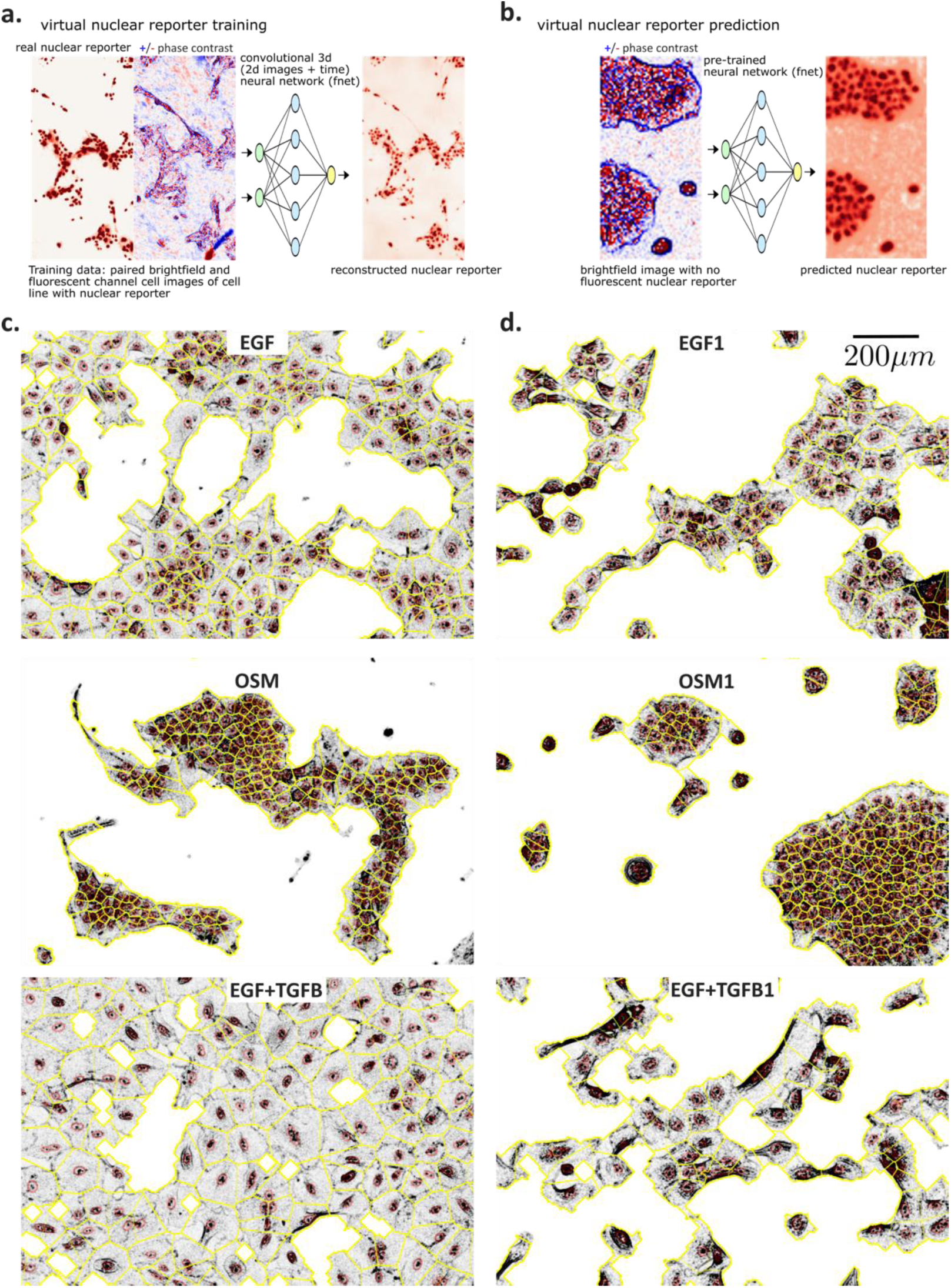
Virtual nuclear staining and nuclear center-based segmentation. a) Paired nuclear reporter (red) and z-normalized phase contrast (+ red - blue) training data input, and reconstructed nuclear reporter output b) Nuclear reporter prediction from out of sample phase contrast images (OSM condition). c) Examples of nuclear segmentations (red) and associated Voronoi boundaries (yellow) overlaid upon phase contrast images (z-normalized absolute value,gray)

**Supplementary Figure 2:**
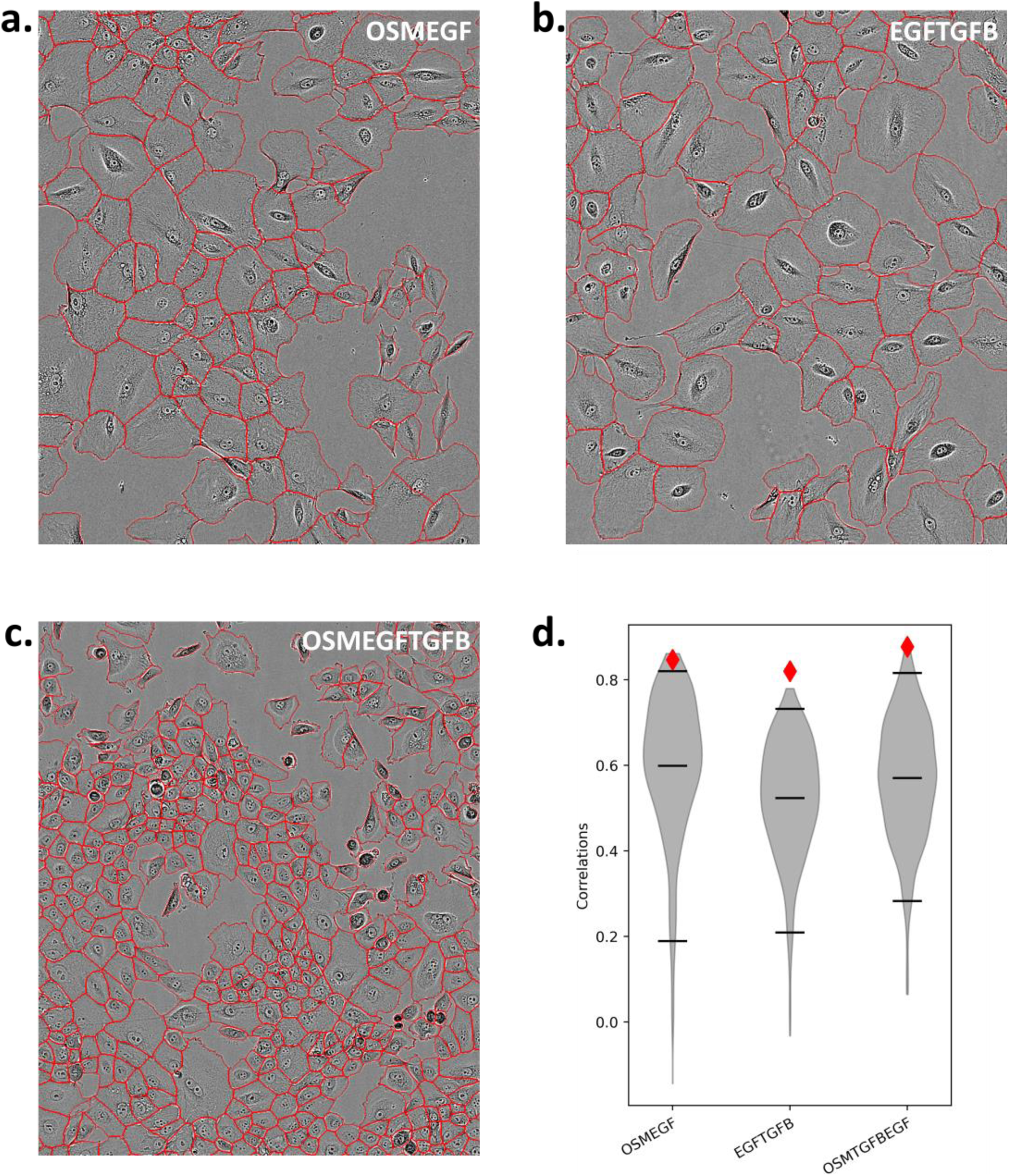
MMIST pipeline applied to a data subset with cytoplasmic masks. Cytoplasmic masks were obtained using the cellpose^115^ software and a subset of the data (only imaging data from the cell-cycle reporter studies. a)-c) Representative images overlaid with cytoplasmic masks from three conditions. Identical tracking and featurization was performed, and the MMIST pipeline was applied to predict RNA-seq at 24 hrs. Predictions were validated via leave-one out cross-validation. d) Model prediction correlation to with-held RNA-seq data from three conditions (red diamonds) and null model (gray violin plots) with the 5^th^ percentile, mean, and 95^th^ percentile of the null distribution indicated (black lines).

**Supplementary Figure 3:**
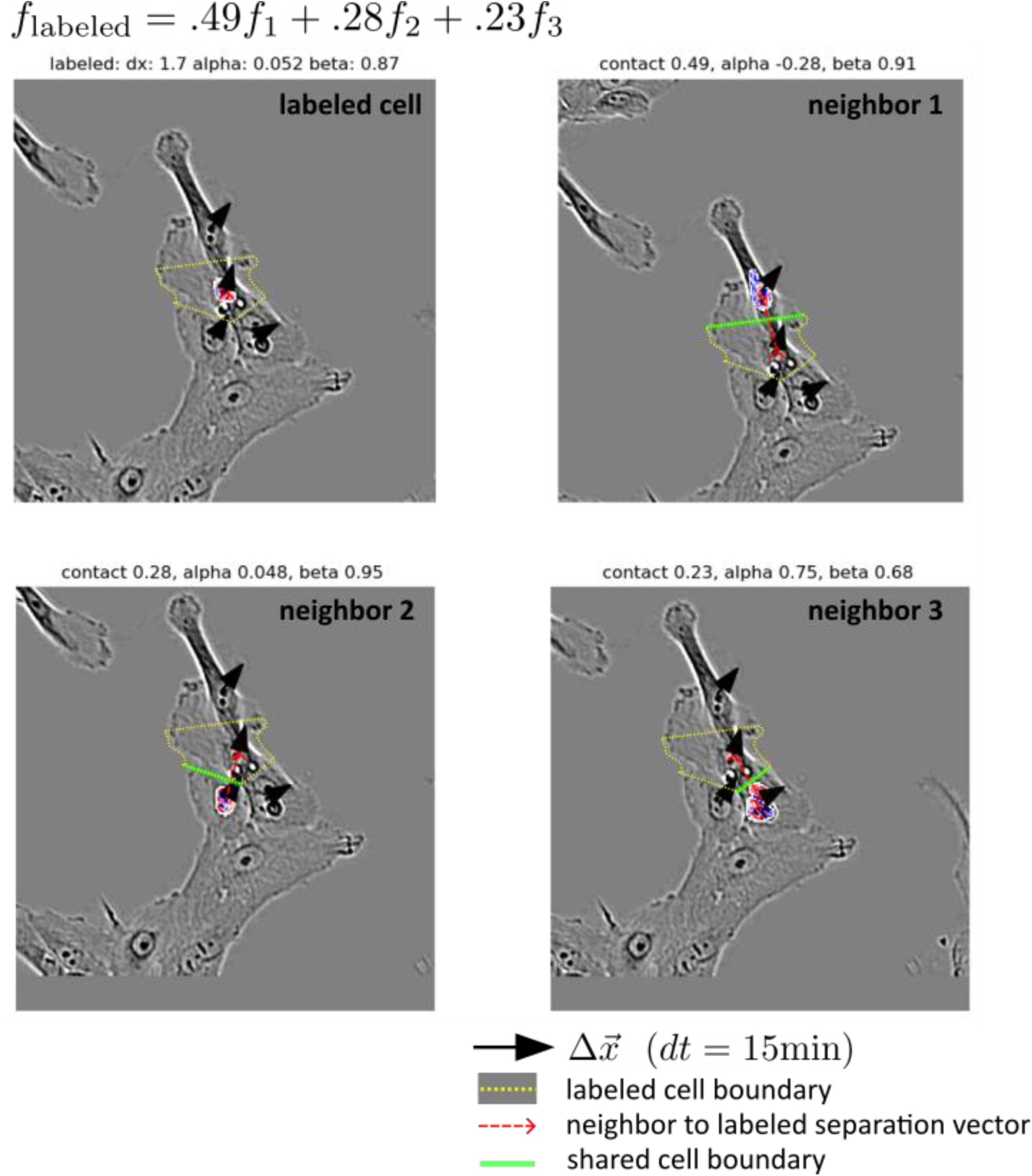
Single-cell and neighborhood motility feature. Single-cell featurization of motility for the labeled cell in the upper right (nuclei in color and Voronoi segmentation in yellow) taken as the magnitude of the displacement from previous frame. The single-cell motility in the context of its local neighborhood taken as the neighbor-weighted average of the 3 boundary cells (upper right, lower left, lower right).

**Supplementary Figure 4:**
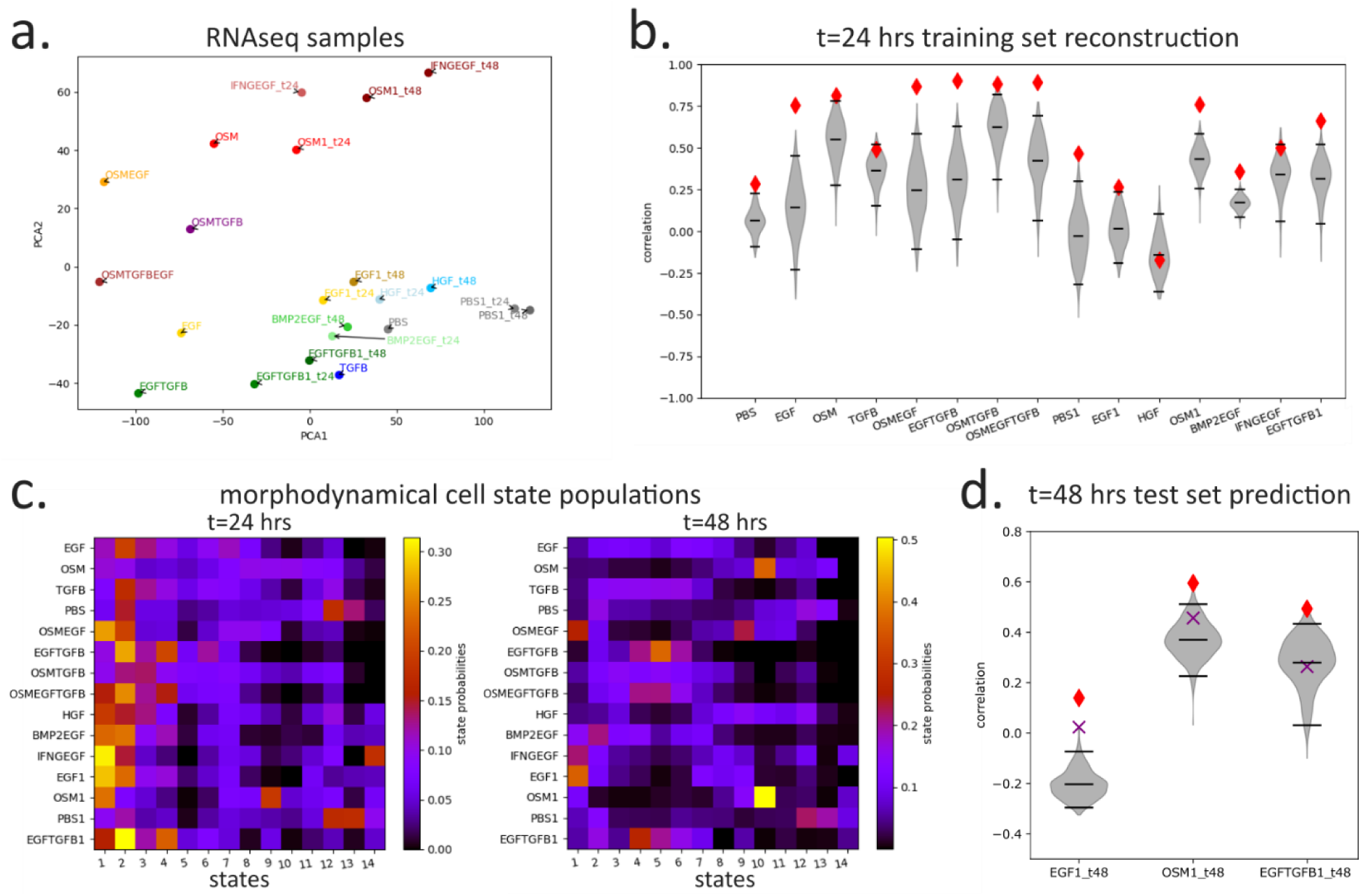
Bulk RNAseq decomposition and time dependence. a) PCA1/2 projection of the RNAseq differential expression, showing sorting by ligand treatment and timepoint. b) Correlation between training set reconstruction and real experimental differential expression. c) Morphodynamical state populations at t=24, 48 hrs d) Pearson correlation to measured RNAseq at 48 hours, MMIST prediction using t=24 hrs trained morphodynamical state gene expression profiles and measured live-cell state populations at t=48hrs (red dots), direct correlation between t=24hr and t=48hr RNAseq (purple x’s), and null with random state populations (gray region).

**Supplementary Figure 5:**
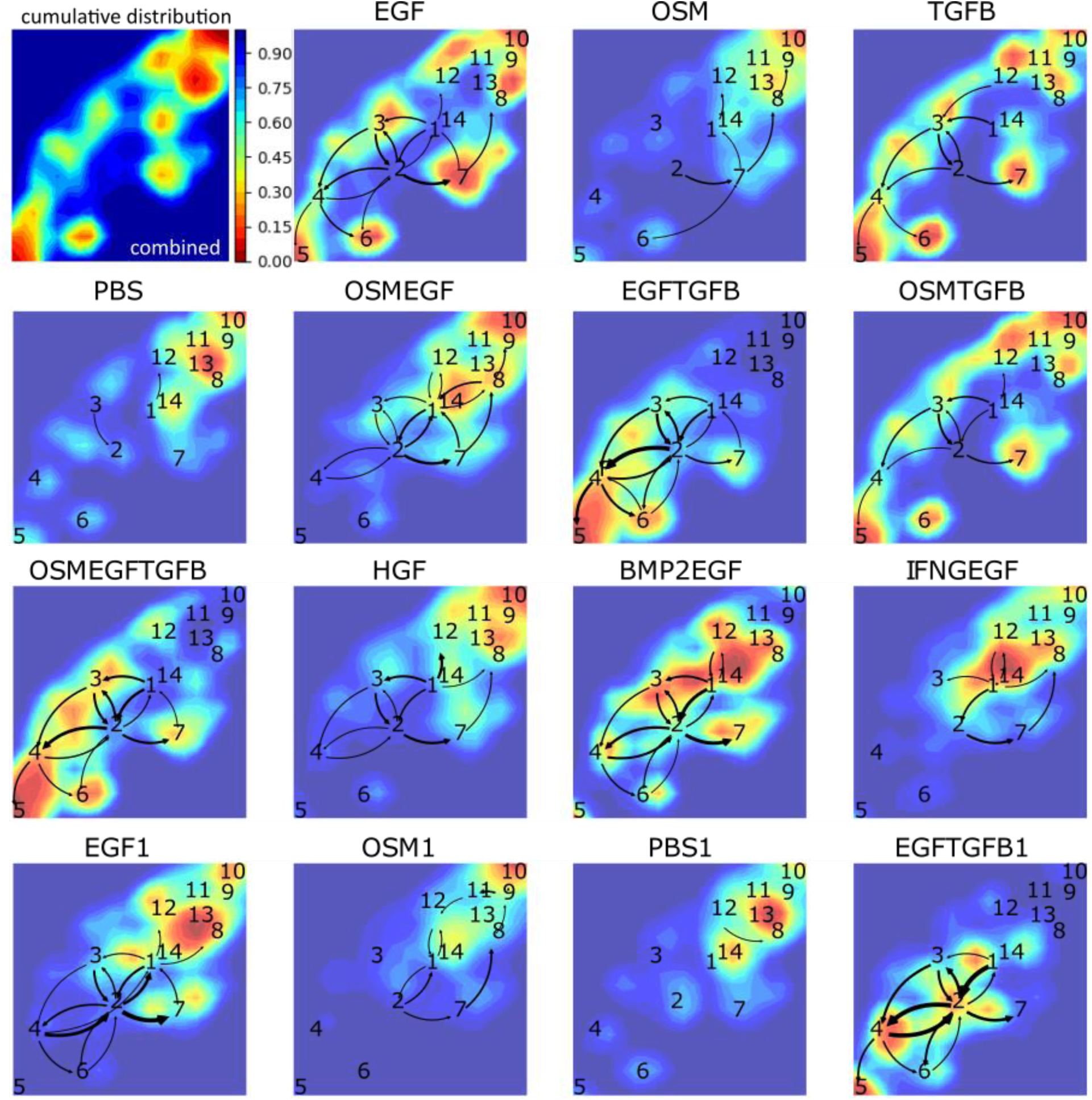
Ligand-dependent populations and cell state flows. Cumulative populations in the UMAP embedding space (blue to red), and state-state transition flows at t=24hrs, in each ligand treatment.

**Supplementary Figure 6:**
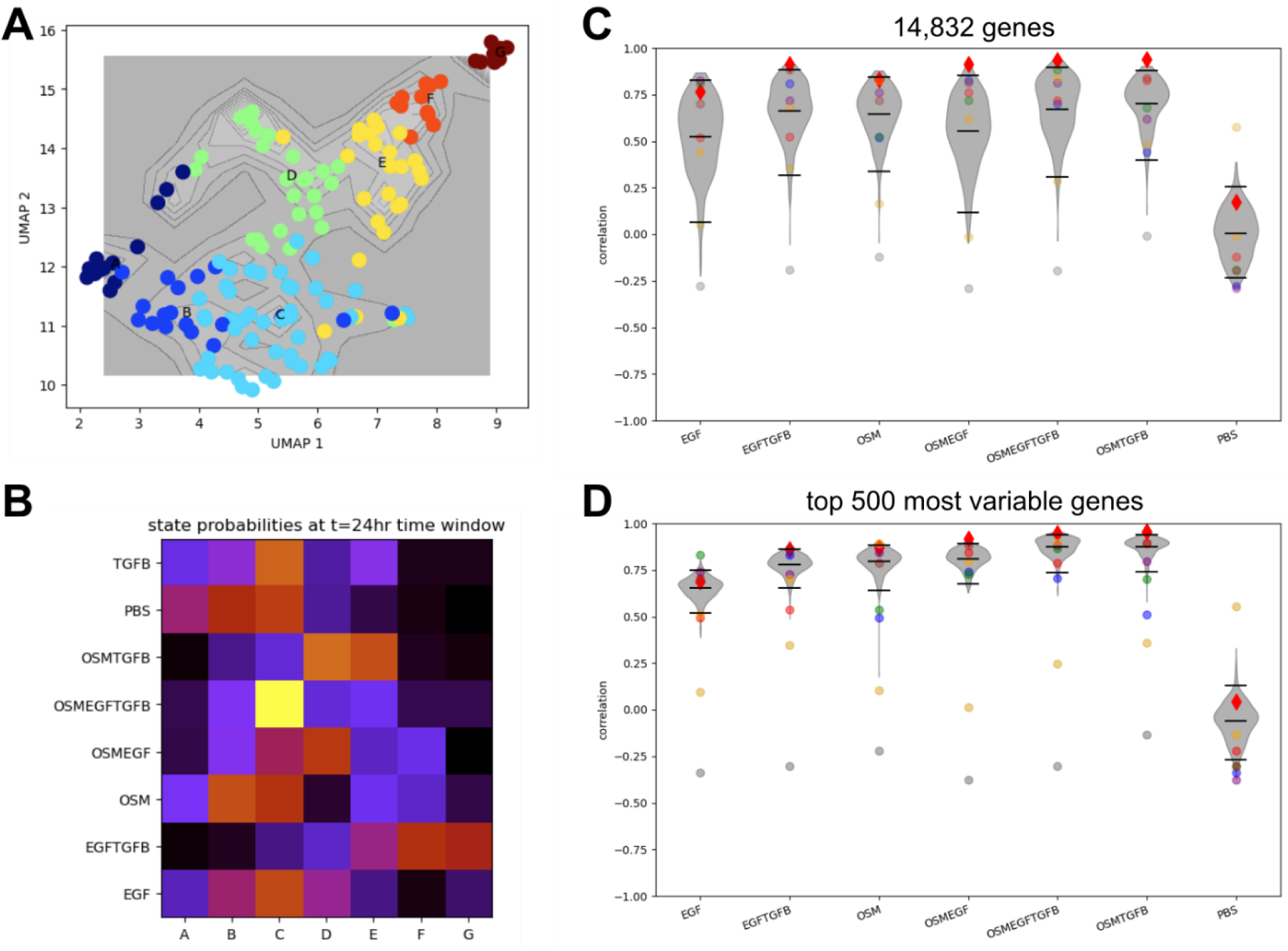
MMIST performance as a function of gene set size. (A) Morphodynamical state populations in a restricted dataset (see https://github.com/jcopperm/celltraj/blob/main/tutorials/mmist.ipynb). (B) Morphodynamical state populations by ligand treatment. (C) MMIST predicted gene expression prediction correlation over all genes in a leave-one-out k-fold cross validation setting (red diamonds), null predictions using random state populations (violin plots with mean, 5^th^ and 95^th^ percentile as black lines), cross treatment correlation between measured ligand treatments (colored circles, EGF:blue, EGFTGFB:green, OSM:red, OSMEGF:purple, OSMEGFTGFB:brown, OSMTGFB:orange, PBS:gray, TGFB:goldenrod). (D)) MMIST predicted gene expression prediction correlation over top 500 most variable genes.

**Supplementary Figure 7:**
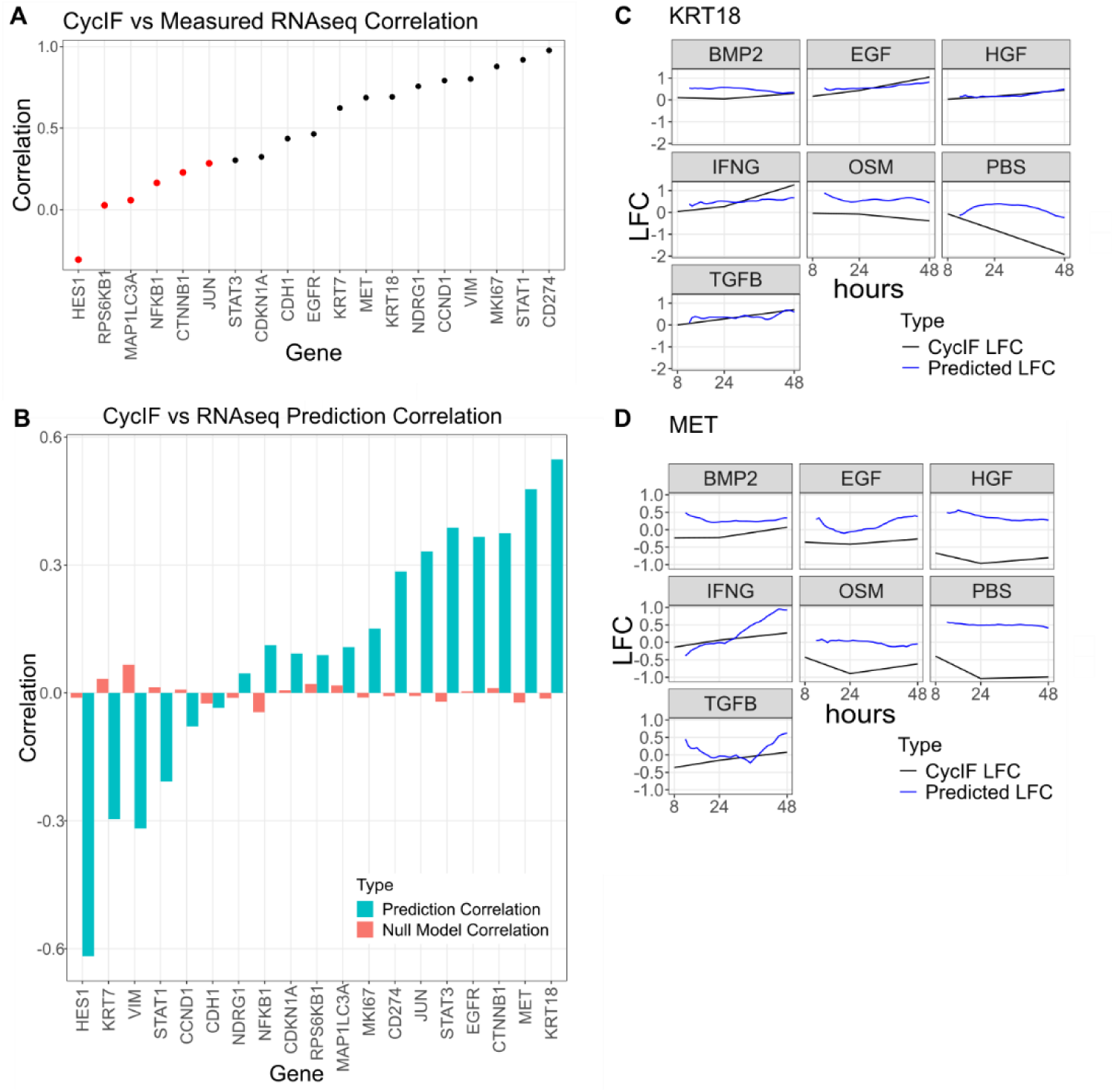
Validation of MMIST prediction of gene expression dynamics from IF images. (A) Overall correlation between population level RNAseq derived RNA levels and cycIF-derived protein levels (black dots considered correlated, red dots uncorrelated). (B) Prediction of timeseries correlation between predicted RNA levels and cycIF derived protein levels by gene averaged over ligand treatment (blue bars) and null prediction with randomly scrambled timepoints. (C,D) Predicted population level gene expression log-fold changes from MMIST (blue lines) and measured protein levels using cycIF (black lines), for the top two best predicted genes.

**Supplementary Figure 8:**
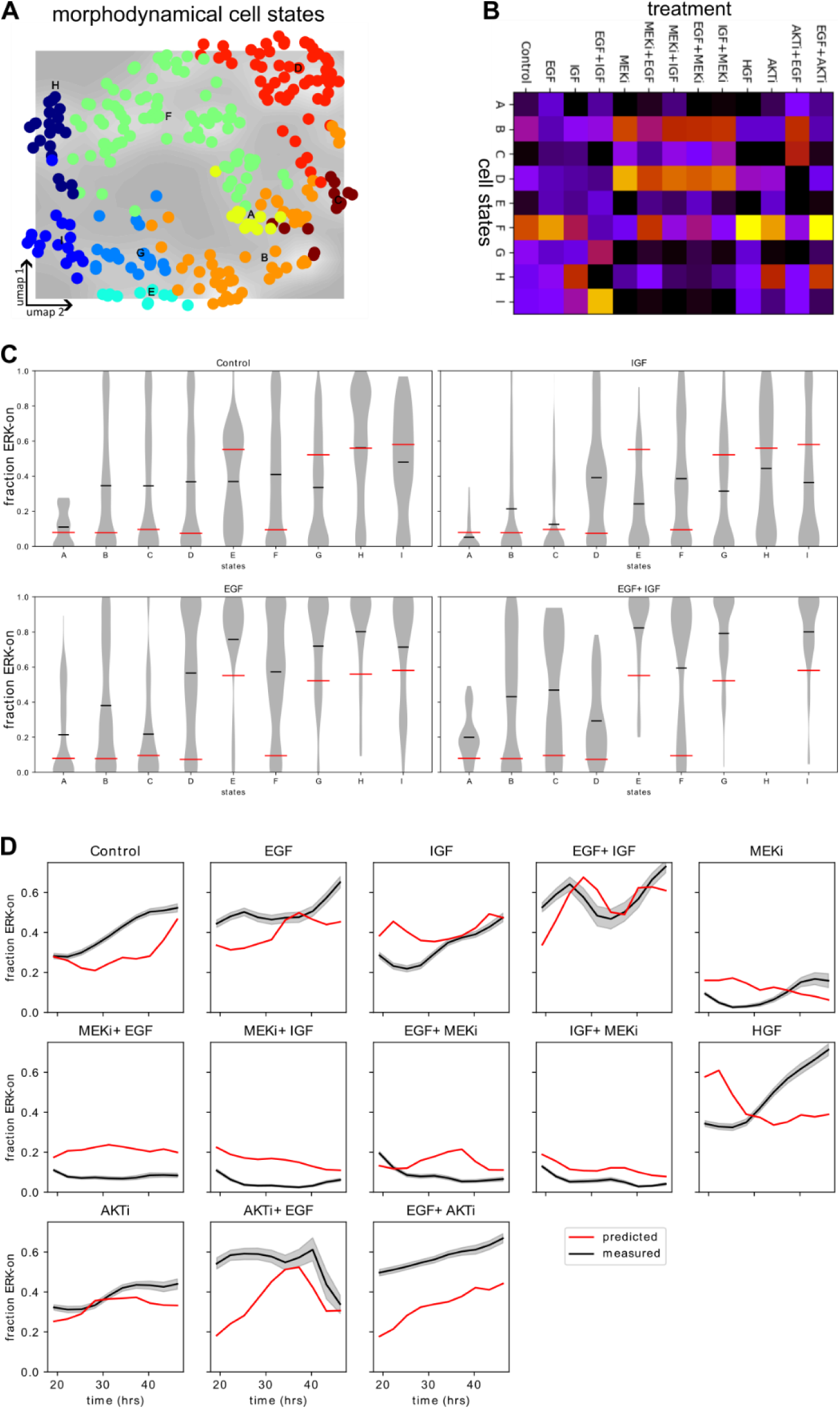
Validation of MMIST prediction of ERK signaling activity dynamics. (A) Morphodynamical cell states derived from 16 hour trajectory embedding (B) State population average (t=24-48hr) by ligand/inhibitor treatment (C) Fraction of ERK “on” (ERKTR c/n ratio >0.8) within morphodynamical cell states (violin plots, mean as black lines) and MMIST derived morphodynamical state levels (red lines). (D) Fraction of ERK “on” cells over time, measured mean and bootstrapped 95% CI (black line and shaded area), and predicted using MMIST (red line).

**Supplementary Figure 9:**
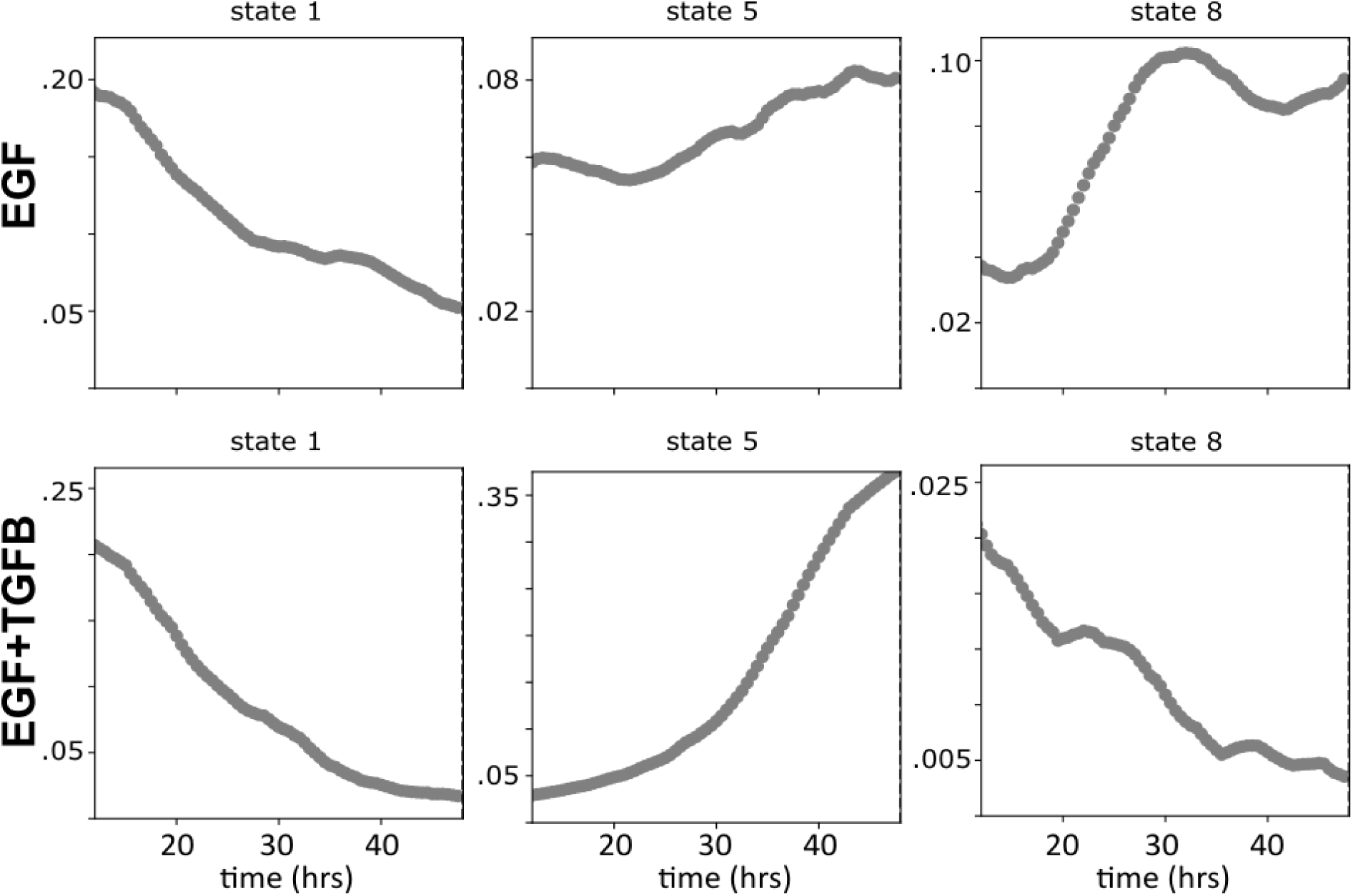
EMT/MET initial and final state probabilities. Live-cell imaging inferred initial and final EMT/MET state populations as a function of time. EMT initial state 1 depopulates over time (leftmost plots), while EMT final state 5 increases over time in EGF+TGFB (lower middle), while MET final state 8 increases over time in EGF conditions (upper right).

**Supplementary Figure 10:**
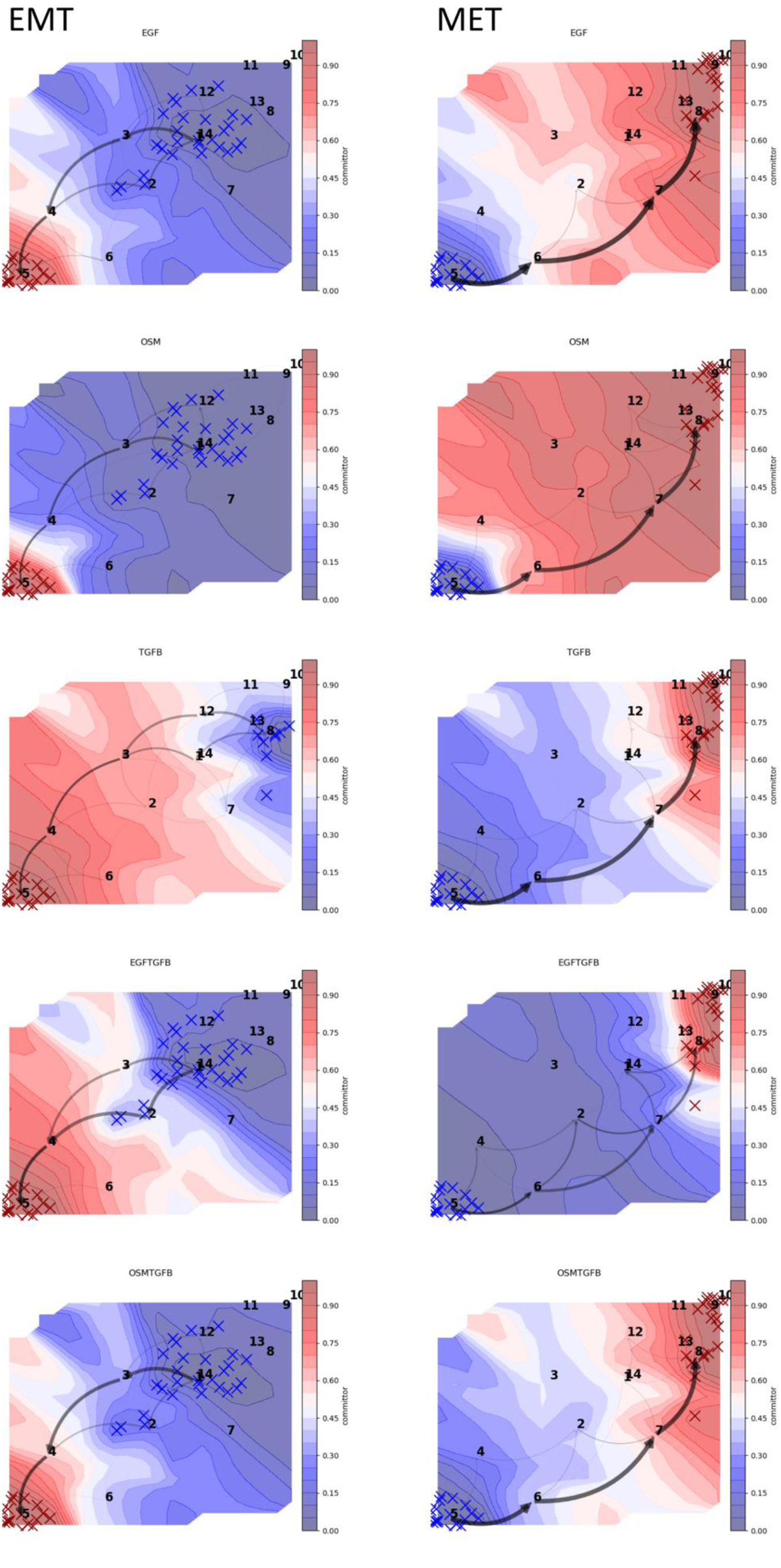
EMT/MET morphodynamical cell state change sequences by ligand treatment. Possible EMT cell state change sequences (initial state 1, final state 5) left, and MET cell state change sequences (initial state 5, final state 8) on the right (black arrows, thickness proportional to transition flux), with final state commitment probability (blue to red) calculated from the 200 k-means state centers and averaged over the UMAP surface.

**Supplementary Figure 11:**
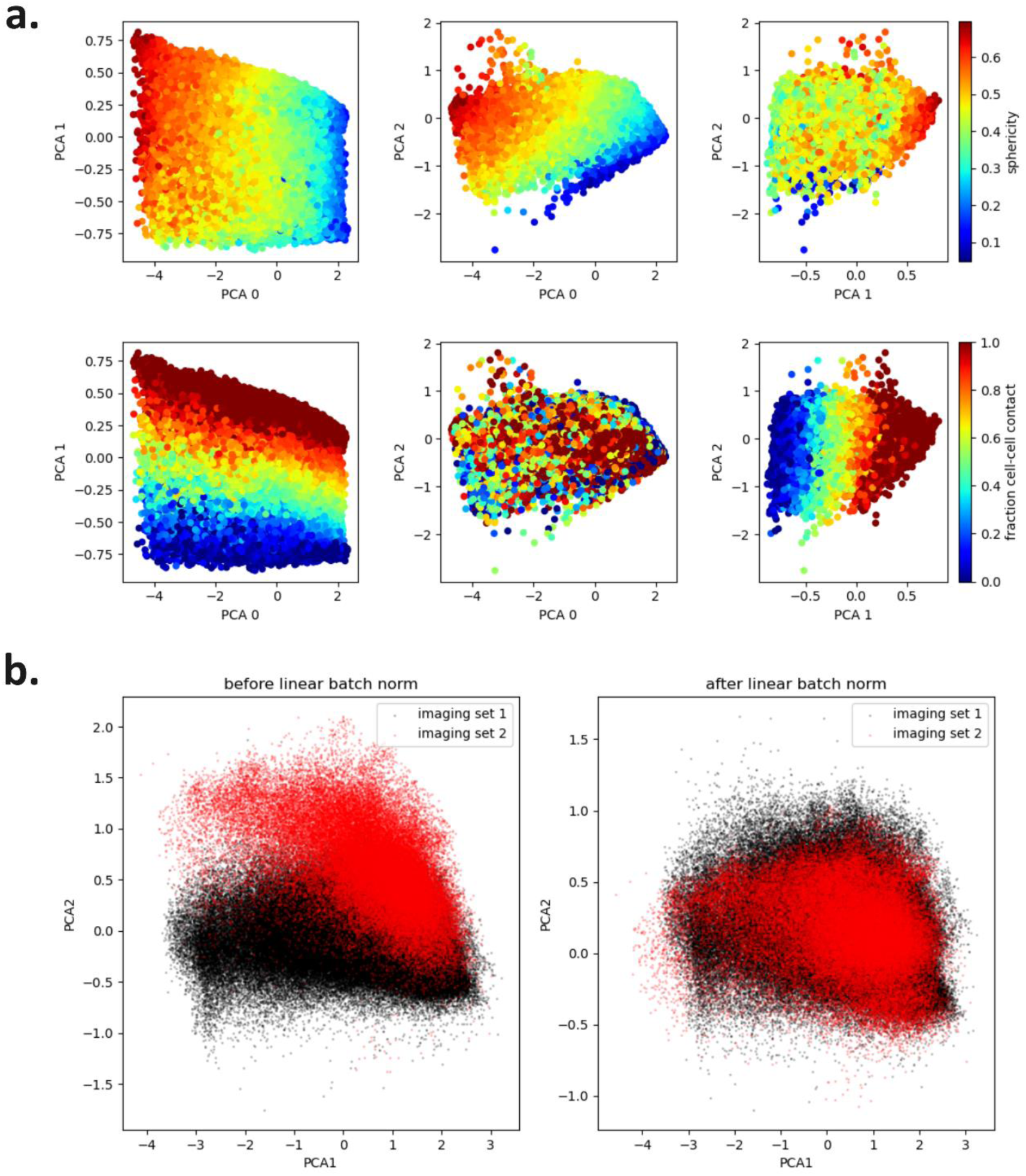
PCA features and feature batch normalization. a.) Scatterplots of top 3 principal components and colored by cell morphology and contact characteristics. b.) Scatterplots of the first two PCA components for imaging experiments 1 (black) and 2 (red), in overlapping treatments, analyzed in this work. Each dot is a cell before (left) and after (right) applying our batch normalization procedure.

**Supplementary Figure 12:**
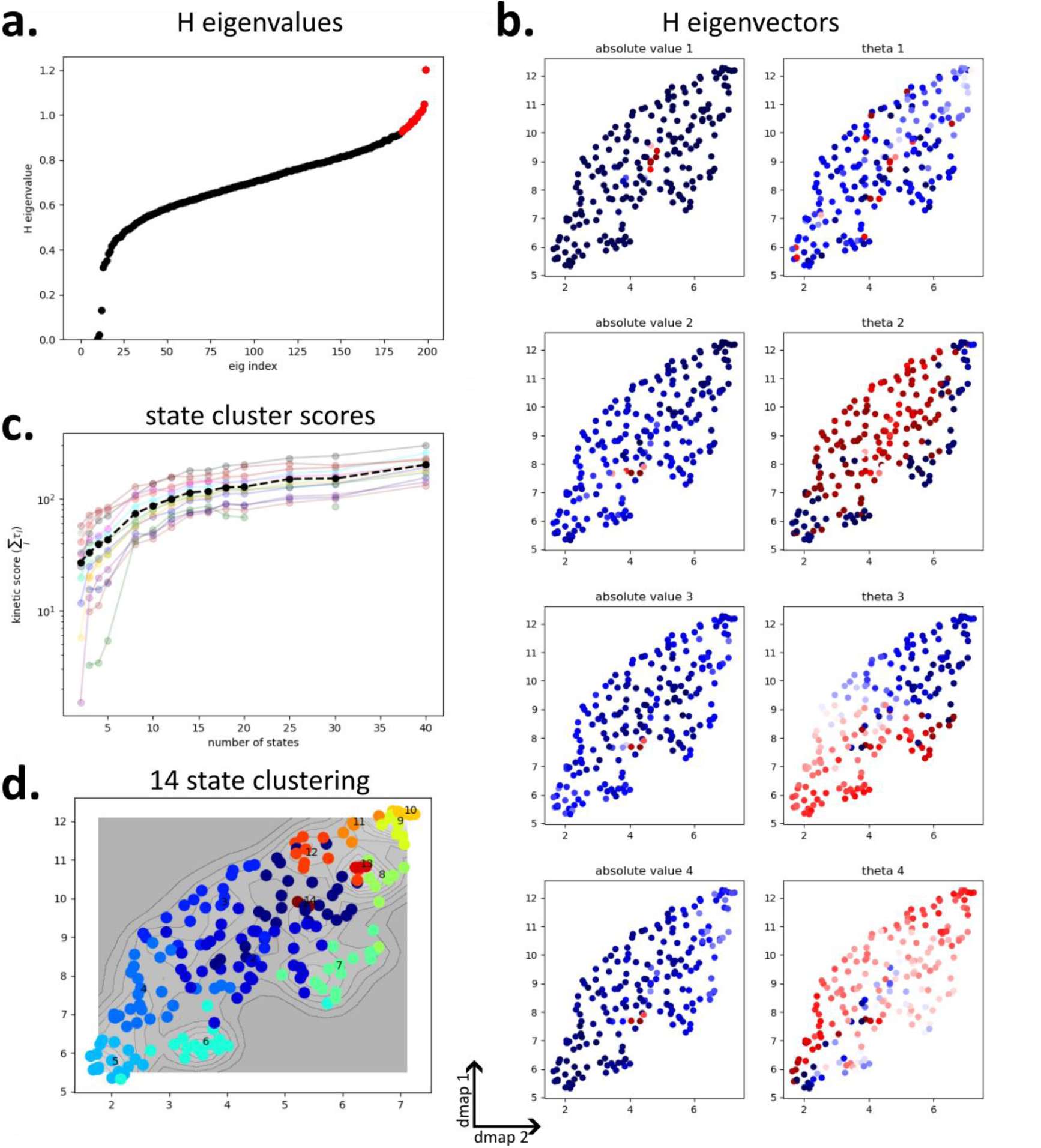
Dynamical clustering of morphodynamical trajectories. (a) Eigenvalues of the Hermitian extension 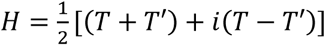 with 𝑇 the transition matrix. (b) 2D UMAP of the eigenvectors of 𝐻 (each point is a microstate of the transition matrix) colored by absolute value and Euler angle of the complex value. (c) State clustering dynamical information quantified by the sum of the timescales from the eigenvalues of 𝑇. The sum of timescales increases rapidly towards 15 states and begins to saturate. (d) K-means clustering into 14 states.

**Supplementary Figure 13:**
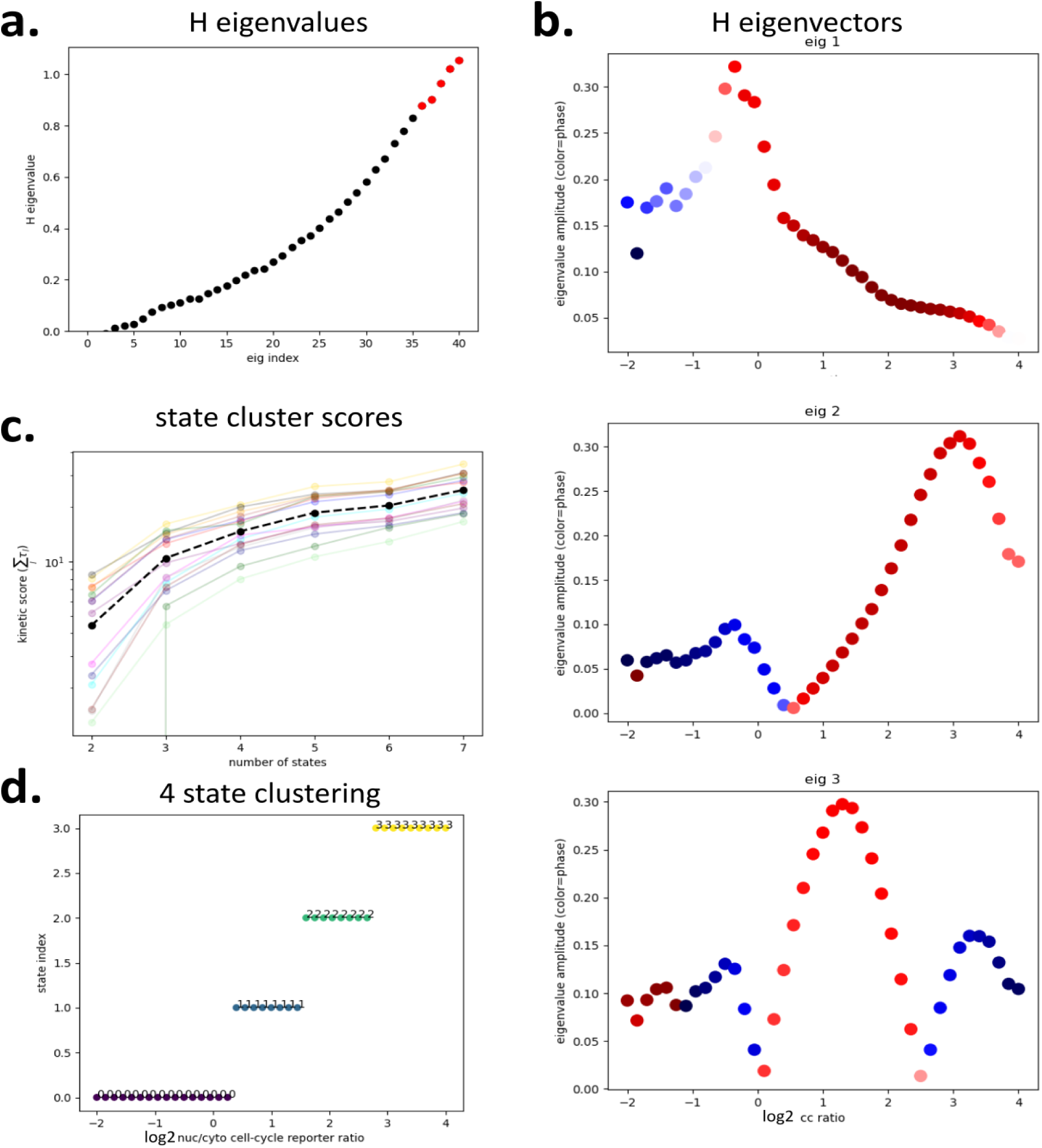
Dynamical clustering of cell-cycle states. (a) Eigenvalues of the Hermitian extension 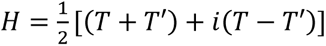 with 𝑇 the transition matrix from dividing log2 of the cell-cycle reporter levels into 51 microstates. (b) Eigenvectors of 𝐻 (each point is a microstate of the transition matrix) colored by Euler angle of the complex value. (c) State clustering dynamical information quantified by the sum of the timescales from the eigenvalues of 𝑇. The sum of timescales increases rapidly towards 4 states and begins to saturate. (d) K-means clustering into 4 states.

**Supplementary Data Table 1.** Segmentation and tracking manual validation. 100 cells per treatment were randomly selected, and evaluated manually to assess segmentation and tracking via the true positive rate (TPR—segmentations or tracks manually validated to be accurate, divided by the total number of cells assessed). Cell identification was assessed by comparing manual counts in a frame at the final timepoint to those obtained from virtually stained nuclear masks.. Segmentation performance from dataset 1 (e.g. EGF1, HGF1) is decreased because for these data the cells did not express a nuclear reporter, and the nucleus was detected via the virtual staining approach only. Tracking performance is decreased as well, due to the decreased segmentation performance and increased time between frames (30 minutes as compared to 15 minutes).

| ligand | PBS | EGF | OSM | TGFB | TGFB<br>+EGF | OSM<br>+EGF | TGFB<br>+OSM | TGFB<br>+OSM<br>+EGF |
| --- | --- | --- | --- | --- | --- | --- | --- | --- |
| count/manual | 99% | 99% | 98% | 96% | 99% | 98% | 96% | 99% |
| seg TPR | 92% | 97% | 97% | 95% | 96% | 96% | 95% | 95% |
| % seg error | 1% | 1% | 1% | 1% | 2% | 2% | 0% | 1% |
| %ambiguous<br>seg | 7% | 2% | 2% | 4% | 2% | 2% | 5% | 4% |
| % tracked | 97% | 91% | 93% | 92% | 95% | 93% | 88% | 90% |
| tracking TPR | 100% | 100% | 98% | 100% | 99% | 99% | 100% | 100% |
| % tracking<br>error | 0% | 0% | 2% | 0% | 0% | 0% | 0% | 0% |
| %ambiguous<br>tracks | 0% | 0% | 0% | 0% | 1% | 1% | 0% | 0% |

| ligand | PBS1 | EGF1 | HGF1 | OSM1 | IFNG+<br>EGF1 | BMP2+<br>EGF1 | EGF+<br>TGFB1 |
| --- | --- | --- | --- | --- | --- | --- | --- |
| count/manual | 79% | 94% | 74% | 90% | 99% | 102% | 113% |
| seg TPR | 80% | 62% | 74% | 86% | 68% | 78% | 75% |
| % seg error | 11% | 25% | 11% | 4% | 24% | 16% | 9% |
| %ambiguous seg | 9% | 10% | 14% | 10% | 8% | 6% | 16% |
| % tracked | 78% | 52% | 78% | 69% | 55% | 66% | 49% |
| tracking TPR | 94% | 94% | 99% | 88% | 91% | 95% | 96% |
| % tracking error | 4% | 6% | 1% | 7% | 5% | 5% | 4% |
| %ambiguous<br>tracks | 2% | 0% | 0% | 5% | 4% | 0% | 0% |

**Supplementary Data Table 2.** Test set gene expression validation with trajectory length.

| test treatment | traj length (hrs) | pred. to exp. corr | diff. from null $\rho - \rho_{\text{null}}$ | upreg true pos. rate | downreg true pos. rate |
| --- | --- | --- | --- | --- | --- |
| OSM+EGF | 0 | .26 | -0.38 | .51 | .80 |
| EGF+TGFB | 0 | .76 | 0.09 | .63 | .89 |
| OSM+EGF+TGFB | 0 | .77 | 0.04 | .77 | .95 |
| OSM+EGF | 1 | .51 | 0.02 | .55 | **.99** |
| EGF+TGFB | 1 | .76 | 0.22 | .68 | .93 |
| OSM+EGF+TGFB | 1 | .75 | 0.14 | .77 | .93 |
| OSM+EGF | 4 | .50 | -0.22 | .51 | .98 |
| EGF+TGFB | 4 | .74 | 0.05 | .68 | .80 |
| OSM+EGF+TGFB | 4 | .73 | -0.04 | .77 | **.96** |
| OSM+EGF | 8 | .55 | -0.15 | .52 | **.99** |
| EGF+TGFB | 8 | .69 | 0 | .68 | **.87** |
| OSM+EGF+TGFB | 8 | .66 | -0.08 | .77 | .94 |
| OSM+EGF | 10 | **.70** | **0.21** | **.72** | .81 |
| EGF+TGFB | 10 | **.81** | **0.27** | **.72** | .84 |
| OSM+EGF+TGFB | 10 | .77 | **0.12** | **.85** | .81 |
| OSM+EGF | 12 | .71 | 0.2 | .70 | .77 |
| EGF+TGFB | 12 | .74 | 0.14 | .69 | .85 |
| OSM+EGF+TGFB | 12 | .64 | 0.01 | .80 | .80 |
| OSM+EGF | 16 | .66 | -0.03 | .65 | .92 |
| EGF+TGFB | 16 | .79 | 0.15 | .70 | .85 |
| OSM+EGF+TGFB | 16 | **.79** | 0.09 | .80 | .88 |

